# Shaped to kill: The evolution of siphonophore tentilla for specialized prey capture in the open ocean

**DOI:** 10.1101/653345

**Authors:** Alejandro Damian-Serrano, Steven H.D. Haddock, Casey W. Dunn

**Affiliations:** Yale University, Department of Ecology and Evolutionary Biology, 165 Prospect St., New Haven, CT 06520, USA; Monterey Bay Aquarium Research Institute, 7700 Sandholdt Rd., Moss Landing, CA 95039, USA

**Keywords:** Siphonophores, nematocysts, predation, specialization, character evolution

## Abstract

Predator specialization has often been considered an evolutionary ‘dead-end’ due to the constraints associated with the evolution of morphological and functional optimizations throughout the organism. However, in some predators, these changes are localized in separate structures dedicated to prey capture. One of the most extreme cases of this modularity can be observed in siphonophores, a clade of pelagic colonial cnidarians that use tentilla (tentacle side branches armed with nematocysts) exclusively for prey capture. Here we study how siphonophore specialists and generalists evolve, and what morphological changes are associated with these transitions. To answer these questions, we: (1) measured 29 morphological characters of tentacles from 45 siphonophore species, (2) mapped these data to a phylogenetic tree, and (3) analyzed the evolutionary associations between morphological characters and prey type data from the literature. Instead of a dead-end, we found that siphonophore specialists can evolve into generalists, and that specialists on one prey type have directly evolved into specialists on other prey types. Our results show that siphonophore tentillum morphology has strong evolutionary associations with prey type, and suggest that shifts between prey types are linked to shifts in the morphology, mode of evolution, and genetic correlations of tentilla and their nematocysts. The evolutionary history of siphonophore specialization helps build a broader perspective on predatory niche diversification via morphological innovation and evolution. These findings contribute to understanding how specialization and morphological evolution have shaped present-day food webs.

**Significance Statement:** Predatory specialization is often associated with the evolution of modifications in the morphology of the prey capture apparatus. Specialization has been considered an evolutionary ‘dead-end’ due to the constraints associated with these morphological changes. However, in predators like siphonophores, armed with modular structures used exclusively for prey capture, this assumption is challenged. Our results show that siphonophores can evolve generalism and new prey-type specializations by modifying the morphological states, modes of evolution, and genetic correlations between the parts of their prey capture apparatus. These findings demonstrate how studying open-ocean non-bilaterian predators can reveal novel patterns and mechanisms in the evolution of specialization. Understanding these evolutionary processes is fundamental to the study of food-web structure and complexity.

## Introduction

Most animal predators use specific structures to capture and subdue prey. Raptors have claws and beaks, snakes have fangs, wasps have stingers, and cnidarians have nematocyst-laden tentacles. The functional morphology of these structures is critical to their ability to successfully capture prey (1). Long-term adaptive evolution in response to the defense mechanisms of the prey (*e*.*g*., avoidance, escape, protective barriers) leads to modifications that can counter those defenses. The more specialized the diet of a predator is, the more specialized its structures need to be to efficiently overcome the challenges posed by the prey. Characterizing the relationships between morphology and predatory specialization is necessary to understand how the phenotypic diversity of predators determines food-web structure. However, for many clades of predators, there is scarce knowledge on how these specializations evolved. The primary questions we set out to answer are: how do predator specialists and generalists evolve, and how does predatory specialization shape morphological evolution?

Siphonophores (Cnidaria: Hydrozoa) are a clade of gelatinous, colonial organisms that swim in the open ocean, feeding on a wide diversity of prey (often fish, crustaceans, and jellyfish). Siphonophores carry modular structures that are exclusively used for prey capture: the tentilla (Fig. 1). The tentilla have great morphological variation across species (2). Together with their well understood function, this makes them an ideal system to study the relationships between functional traits and prey specialization. Like a head of coral, a siphonophore is a colony bearing many feeding polyps (Fig. 1). Each feeding polyp has a single tentacle, which branches into a series of tentilla (side branches). Like other cnidarians, siphonophores capture prey with nematocysts, harpoon-like stinging capsules borne within specialized cells known as cnidocytes. Unlike the prey capture apparatus of most other cnidarians, siphonophore tentacles carry their cnidocytes in extremely complex and organized batteries (3) which are located in their tentilla. While nematocyst batteries and clusters in other cnidarians are simple static scaffolds for cnidocytes, siphonophore tentilla have their own reaction mechanism, triggered upon encounter with prey. When it fires, a tentillum undergoes an extremely fast conformational change that wraps it around the prey, maximizing the surface area of contact for nematocysts to fire on the prey (4). In addition, some species have elaborate fluorescent and bioluminescent lures on their tentilla to attract prey with aggressive mimicry (5–7).

**Figure 1:**
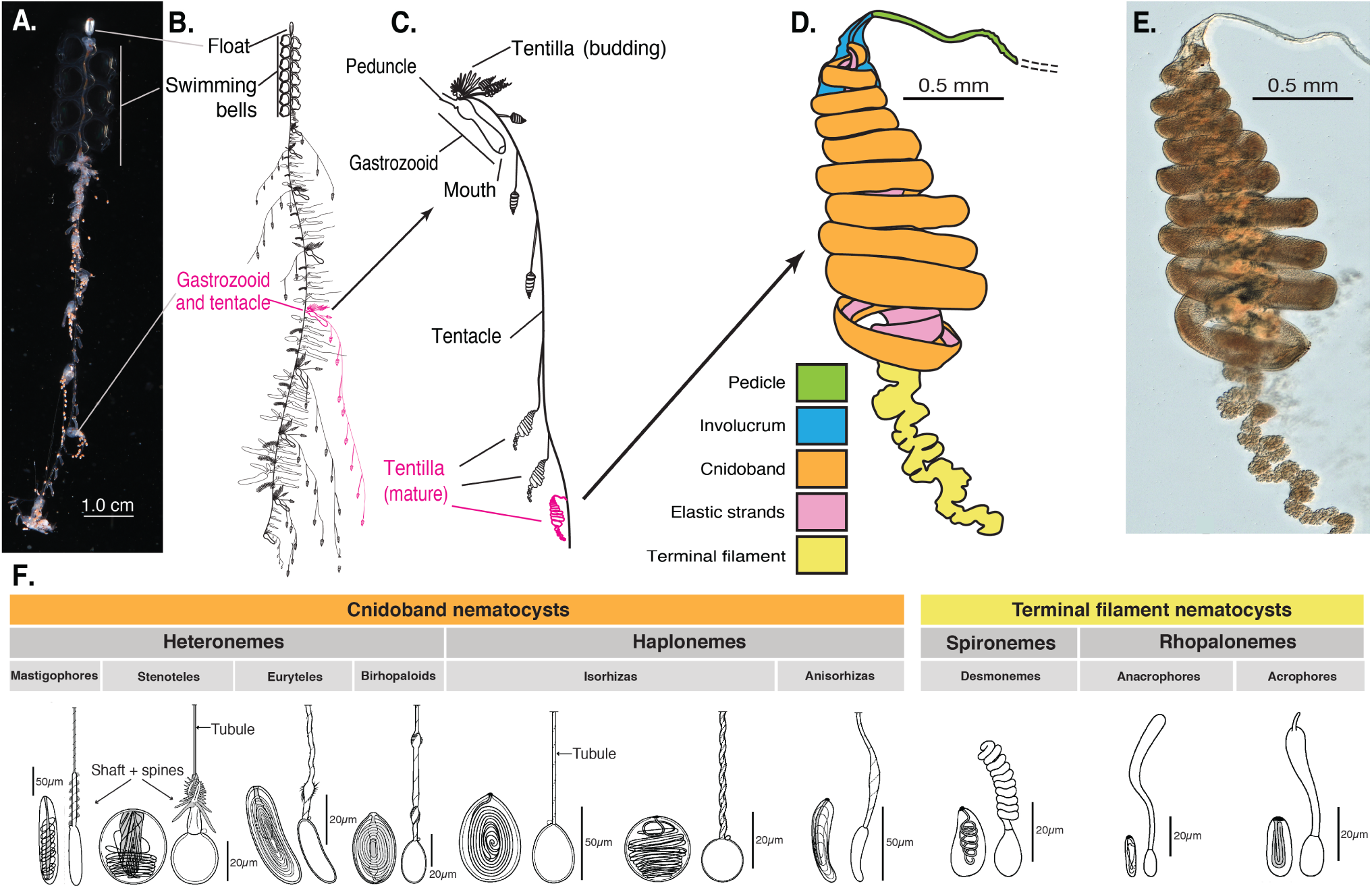
Siphonophore anatomy. A - *Nanomia* sp. siphonophore colony (photo by Catriona Munro). B, C - Illustration of a *Nanomia* colony, gastrozooid, and tentacle closeup (by Freya Goetz). D - *Nanomia* sp. Tentillum illustration and main parts. E - Differential interference contrast micrograph of the tentillum illustrated in D. F - Nematocyst types (illustration reproduced with permission from Mapstone 2014), hypothesized homologies, and locations in the tentillum. Undischarged to the left, discharged to the right.

Siphonophores bear four major nematocyst types in their tentacles and tentilla (Fig. 1F).. The largest type, heteronemes, have open-tip tubules characterized by bearing a distinctly wider spiny shaft at the proximal end of the everted tubule. These are typically found flanking the proximal end of the cnidoband. The most abundant type, haplonemes, have no distinct shaft, but similarly to heteronemes, their tubules have open tips and can be found in the cnidoband. Both heteronemes and haplonemes bear short spines along the tubule. Both can be toxic and penetrate the surface of some prey types. In the terminal filament, siphonophores bear two other types of nematocysts, characterized by their adhesive function, closed tip tubules, and lack of spines on the tubule. These are the desmonemes (a type of adhesive coiled-tubule spironeme), and rhopalonemes (a siphonophore-exclusive nematocyst type with wide tubules).

Many siphonophore species inhabit the deep pelagic ocean, which spans from ∼200m to the abyssal seafloor (∼4000m). This habitat has fairly homogeneous physical conditions and stable zooplankton abundances and composition (8). With relatively predictable prey availability, ecological theory predicts that interspecific competition would inhibit the coexistence of closely-related species unless evolution towards specialization reduces the breadth of each species’ niche (9–11). If this prediction holds true, we would expect the prey-capture apparatus morphologies of siphonophores to diversify with the evolution of specializations on a variety of prey types in different siphonophore lineages.

Specialization has been thought to be an evolutionary ‘dead-end’, meaning that specialized lineages are unlikely to evolve into generalists or to shift the resource for which they are specialized (12–16). However, recent studies have found that interspecific competition can favor the evolution of generalists from specialists (17–19) and specialist resource switching (20, 21). In addition to studying relationships with morphology, we seek to identify what evolutionary transitions in trophic niche breadth are prevalent in these open-ocean tactile predators. To do so, we examine three alternative scenarios of siphonophore trophic specialization: (1) predatory specialists evolved from generalist ancestors; (2) predatory specialists evolved from specialist ancestors which targeted different resources, switching their primary prey type; and (3) predatory generalists evolved from specialist ancestors. These scenarios are non-exclusive, and each could apply to different transitions along the siphonophore phylogeny.

In the past, the study of siphonophore tentilla and diets has been limited due to the inaccessibility of their oceanic habitat and the difficulties associated with the collection of fragile siphonophores. Thus, the morphological diversity of tentilla has only been characterized for a few taxa, and their evolutionary history remains largely unexplored. Contemporary underwater sampling technology provides an unprecedented opportunity to explore the trophic ecology (22) and functional morphology (23) of siphonophores. In addition, well-supported phylogenies based on molecular data are now available for these organisms (24). These advances allow for the examination of the evolutionary relationships between modern siphonophore form, function, and ecology.Our work builds upon previous pioneering studies that have explored the relationships between tentilla and diet, and showed that siphonophores are a robust system for the study of predatory specialization via morphological diversification. Purcell (25, 26) showed clear relationships between diet, tentillum, and nematocyst characters in co-occurring epipelagic siphonophores for a small subset of extant epipelagic siphonophore species.

In this study, we present an extensive morphological characterization of tentilla and their nematocysts across a broad variety of shallow and deep-sea siphonophore species using modern imaging technologies, summarize the literature on siphonophore diets, expand the phylogenetic tree of siphonophores by combining ribosomal gene sequences from a broad range of taxa with a transcriptome-based backbone tree, and explore the evolutionary histories and correlations between diet, tentillum, and nematocyst characters. Our results suggest that siphonophores can evolve new specializations and generalism by modifying the phenotypes and genetic correlations in their prey capture apparatus. These findings show how studying elusive non-bilaterian predators can challenge traditional views on the evolution of predatory specialization.

## Results

### Novel phylogenetic relationships

In order to analyze the relationships between morphology and diet across the evolutionary history of siphonophores, we generated a siphonophore phylogeny that had broader taxonomic sampling than was available in previously published analyses. We first inferred a new tree with the needed taxon sampling with publicly available ribosomal RNA genes (18S & 16S) and new data from one species. This tree is essentially an extended version of that published in (27), and the two are congruent. We then compared the new extended ribosomal RNA tree to a recently published siphonophore transcriptome phylogeny (24). The topology of the extended ribosomal RNA tree recapitulates the resolved nodes in (27) and most of the nodes in (24). Only five nodes in the unconstrained tree inference were incongruent with the (24) transcriptome tree, with four of them poorly supported (bootstrap values <84), and only one of them strongly supported (*Frillagalma vityazi*-*Nanomia bijuga*, 100 bootstrap support). We constrained the incongruent nodes to the (24) topology during estimation of the constrained 18S+16S tree inference (Fig. 2). Since the transcriptome-based placement of *Nanomia bijuga* is more consistent with the morphological data, that relationship was also constrained. Moreover, with the inclusion of sequences from *Stephanomia amphytridis* and multiple *Erenna* species, our tree reveals a novel sister relationship between the genus *Erenna* and *Stephanomia*.

**Figure 2:**
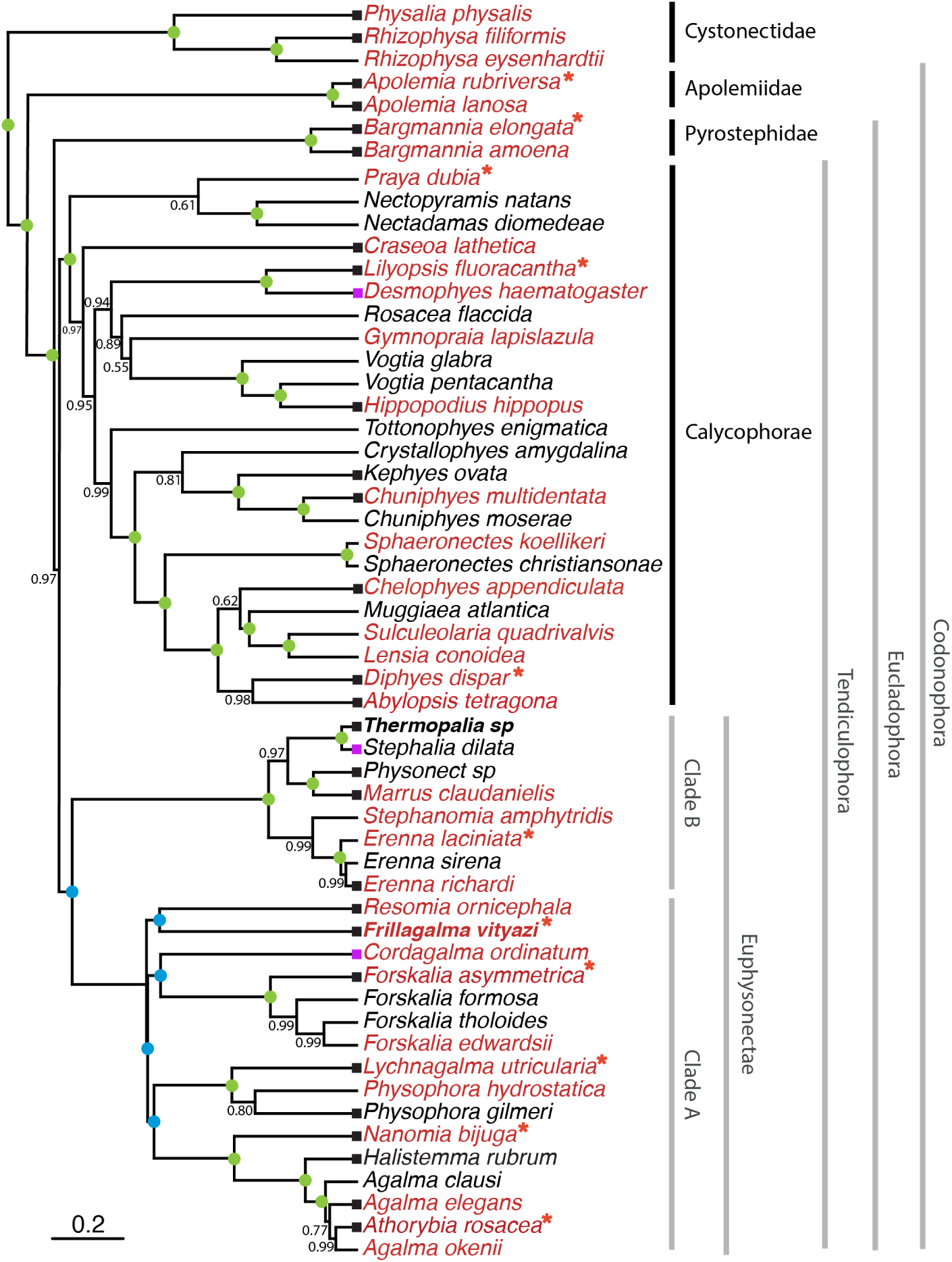
Bayesian time-tree inferred from 18S + 16S concatenated sequences and constrained to be congruent with a published transcriptome phylogeny. Branch lengths estimated using a relaxed molecular clock. Species names in red indicate replicated representation in the morphology data. All data were publicly available, apart from new sequences produced for Thermopalia taraxaca and Frillaglma vityazi. Nodes labeled with Bayesian posteriors (BP). Green circles indicate BP = 1. Blue circles indicate nodes constrained to be congruent with Munro *et al*. (2018). Tips with black squares indicate the species with transcriptomes used in Munro *et al*. (2018). Tips with purple squares indicate genus-level correspondence to taxa included in Munro *et al*. (2018). The main clades are labeled: with black bars for described taxonomic units, and grey bars for operational phylogenetic designations.

We used the clade nomenclature defined in (27) and (24), including Codonophora to indicate the sister group to Cystonectae, Euphysonectae to indicate the sister group to Calycophorae, Clade A and B to indicate the two main lineages within Euphysonectae. In addition, we define two new clades within Codonophora (Fig. 2): Eucladophora as the clade containing *Agalma elegans* and all taxa that are more closely related to it than to *Apolemia lanosa*, and Tendiculophora as the clade containing *Agalma elegans* and all taxa more closely related to it than to *Bargmannia elongata*. Eucladophora is characterized by bearing spatially differentiated tentilla with proximal heteronemes and a narrower terminal filament region. The etymology derives from the Greek *eu*+*kládos*+*phóros* for “true branch bearers”. Tendiculophora are characterized by bearing rhopalonemes and desmonemes in the terminal filament, having a pair of elastic strands, and developing proximally detachable cnidobands. The etymology of this clade is derived from the Latin *tendicula* for “snare or noose” and the Greek *phóros* for “carriers”.

### Evolutionary associations between diet and tentillum morphology

We reconstructed the evolutionary history of feeding guilds using stochastic mapping on the new phylogeny (Fig. 3). Our reconstructions do not recover generalism as the ancestral siphonophore diet. None of the transitions in diet are consistent with scenario 1 (specialists evolving from generalists). Feeding guild specializations have shifted from an alternative ancestral state at least five times, consistent with instances supporting scenario 2 (specialists evolving to feed on a different resource). We also recover multiple independent origins of generalism from specialist ancestors (Fig. 3). Large crustacean specialists evolve into generalists twice independently, consistent with instances of scenario 3 (generalists evolving from specialists). This finding is particularly compelling given in that it is the opposite of known biases in ancestral state reconstruction. (28) found that such methods tend to infer higher transition rates toward the more frequent state. In this case, that would lead to a bias for an increased rate of transition from generalists (the rarer state across the tips) to specialists (the more common state across the tips). We observe the opposite, indicating strong evidence that these generalists are indeed a derived state.

**Figure 3:**
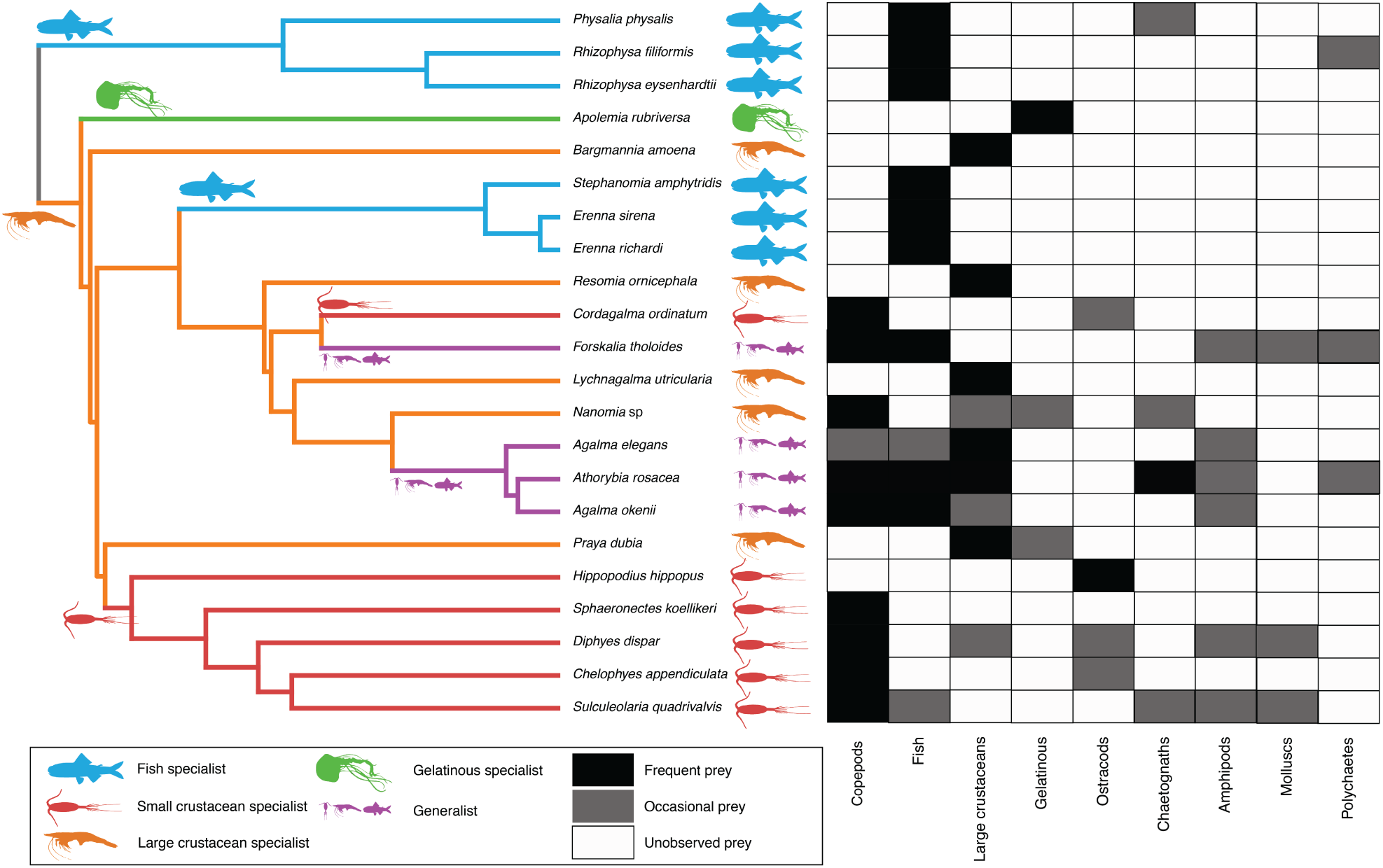
Left - Subset phylogeny showing the mapped feeding guild regimes that were used to inform the *OUwie* analyses. Right - Grid showing the prey items consumed from which the feeding guild categories were derived. Diet data were obtained from the literature review, available in the Dryad repository.

To test whether measured morphological characters evolved in association with shifts in feeding ecology, we analyzed the evolutionary history of each character on the phylogeny, with the feeding guilds reconstructed on it as hypothetical selective regimes. We fit and compared alternative evolutionary models for each continuous character. The models compared were the Brownian Motion (BM) model of neutral divergent evolution (29), the Ornstein-Uhlenbeck (OU) model of stabilizing selection around a single fitted optimum state (30, 31), and an OU model with multiple optima (OUm) corresponding to each reconstructed selective regime (feeding guild). The model comparison shows that out of 30 characters, 10 show significantly stronger support for the diet-driven multi-optima OU model (S15). These characters include terminal filament nematocyst size and shape, involucrum length, elastic strand width, and heteroneme number. Most of these characters are found exclusively in Tendiculophora, thus this may reflect processes that could be unique to this subtree. Five characters including cnidoband length, cnidoband shape, and haploneme length show maximal support for a diet-driven single-optimum OU model. The remaining 15 characters support BM (or OU with marginal AICc difference with BM).

In order to investigate the associations between the evolutionary history of morphological characters and specific prey types found in the diet, we used phylogenetic logistic regressions. We found that several characters were significantly correlated with the gains and losses of specific prey types (Fig. 3, right). Shifts toward ostracod presence in diet correlated with reductions in pedicle width and total haploneme volume. Shifts to copepod presence in the diet were associated with reductions in haploneme width, cnidoband length and width, total haploneme and heteroneme volumes, and tentacle and pedicle widths. Consistently, transitions to decapod presence in the diet correlated with more coiled cnidobands (S21). Evolutionary shifts in these characters may have allowed the inclusion of these prey types in the diet.

In addition to studying correlations with prey type presence/absence in the diet, we also tested for correlations between morphological characters and shifts in prey selectivity using phylogenetic linear models. Prey selectivity values were calculated from (32) by contrasting the gut content frequencies to the corresponding environmental abundances of prey. We found that fish selectivity is associated with increased number of heteronemes per tentillum, increased roundness of nematocysts (desmonemes and haplonemes), larger heteronemes, reduced heteroneme/cnidoband length ratios, smaller rhopalonemes, lower haploneme surface area to volume ratio (SA/V), and larger the cnidoband, elastic strand, pedicle and tentacle widths. Decapod-selective diets were associated with increasing cnidoband size and coiledness, haploneme row number, elastic strand width, and heteroneme number. Copepod-selective diets evolved in association with smaller heteroneme and total nematocyst volumes, smaller cnidobands, rounder rhopalonemes, elongated heteronemes, narrower haplonemes with higher SA/V ratios, and smaller heteronemes, tentacles, pedicles, and elastic strands. Selectivity for ostracods was associated with reductions in size and number of heteroneme nematocysts, cnidoband size, number of haploneme rows, heteroneme numbers, and cnidoband coiledness. Heteroneme length and elongation also correlated negatively with chaetognath selectivity (S21). These results indicate that not only diet but also differential feeding selectivity has evolved in correlation with changes in the prey capture apparatus of siphonophores. For each prey type studied, tentillum morphology is a much better predictor of prey selectivity than of prey presence in the diet, despite prey selectivity data being available for a smaller subset of species. Interestingly, many of the morphological predictors had opposite slope signs when predicting prey selectivity *versus* predicting prey presence in the diet (Table 2).

**Table 1.** Tests of correlated evolution between siphonophore morphological characters and aspects of the diet found correlated in the literature. We report the direction and significance of the evolutionary association, the number of taxa used for the analysis, and the literature source where the morphology-diet association was first reported.

| Character | Aspect of diet | Test of evolutionary association | Relationship sign | P-value | Number of taxa | Association first report |
| --- | --- | --- | --- | --- | --- | --- |
| Differentiated cnidobands | Hard bodied prey | Pagel's test | + | 0.017 | 19 | Purcell, 1984 |
| Heteroneme volume | Copepod prey size | pGLS | + | 0.002 | 8 | Purcell, 1984 |
| Terminal filament nematocysts | Crustacean diet | Pagel's test | Non-Significant | 0.200 | 19 | Purcell & Mills, 1988 |
| Number of nematocyst types | Soft-bodied prey | Phylogenetic logistic regression | - | 0.040 | 22 | Purcell & Mills, 1988 |

**Table 2.**
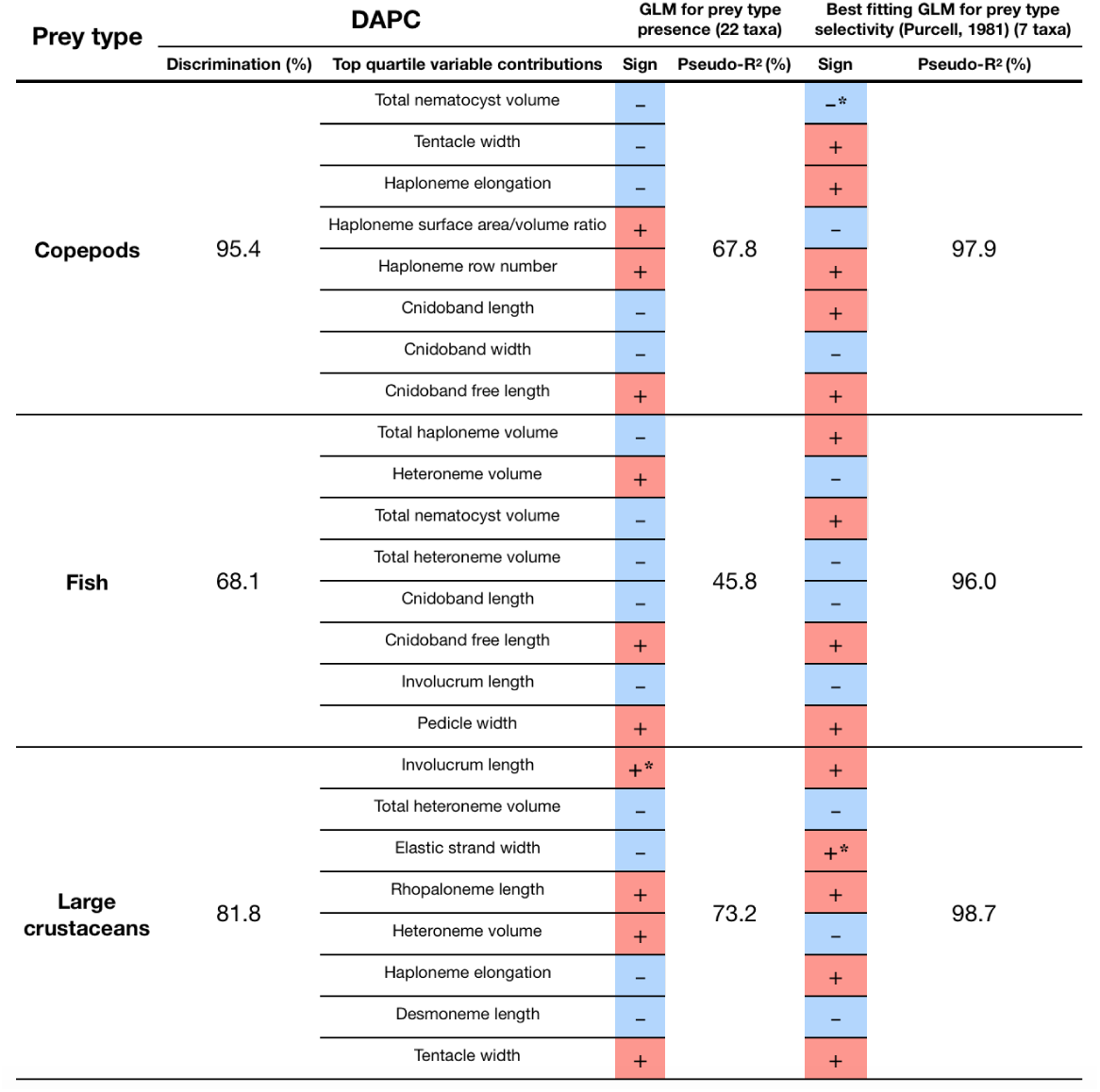
Discriminant analysis of principal components for the presence of specific prey types using the morphological data. Top quartile variable (character) contributions to the linear discriminants are ordered from highest to lowest. Logistic regressions and GLMs were fitted to predict prey type presence and selectivity respectively. The sign of the slope of each predictor is reported, marked with an asterisk if significant (p-value < 0.05), and highlighted grey if it differs between prey presence in diet and prey selectivity. Pseudo-R^2^ (%) approximates the percent variance explained by the model.

We tested some of the diet-morphology associations previously proposed in the literature (25, 26) for correlated evolution (Table 1). We found that most, such as heteroneme volume and copepod prey size, do show evidence for correlated evolution. The sole exception was the relationship between terminal filament nematocysts (rhopalonemes and desmonemes) and crustaceans in the diet. Analyses that do not take phylogeny into account do recover this correlation across the extant species studied, but it is not consistent with correlated evolution. The latter is likely a product of the larger species richness of crustacean-eating species with terminal filament nematocysts, rather than simultaneous evolutionary gains.

### Evolution of relationships between characters with diet

Phenotypic integration results in correlation patterns between morphological characters and their rates of evolution. To study these patterns, we fit a set of evolutionary variance-covariance matrices (33). The quantitative characters we measured from tentilla and their nematocysts are highly correlated. The results indicate that the dimensionality (number of independent axes of variation) of tentillum morphology is low, that many traits are associated with size, but that nematocyst arrangement and shape are independent of it (S4). The variance-covariance matrices (S21-23) are congruent with the abundant positive correlations observed among simple measurement characters in S3. This analysis more clearly reveals the diagonal blocks that constitute the evolutionary modules, such as the heteroneme block, the terminal filament nematocyst block, and the cnidoband-pedicle-tentacle block. These results were not sensitive to the transformation of inapplicable states and taxon sampling. These results indicate that siphonophore tentilla and nematocysts are phenotypically integrated and co-evolve within discrete evolutionary modules.

In order to test whether rate covariance matrices changed with evolutionary shifts in feeding guild regimes, we compared the rate covariance terms between characters across the subtrees occupied by the different feeding guild regimes (S21). We found that half (48%) of the character pairs presented significantly distinct correlation coefficients across different regimes (S19), indicating that the mode of phenotypic integration also shifts with trophic niche. When contrasting the regime-specific rate correlation matrices to the whole-tree matrix and to the preceding ancestral regime matrix, we were able to identify the character dependencies that are unique to each predatory niche (S22-23).

We were able to identify specific character correlations that shifted with the evolution of new diets. Under the majority of stochastic character mapping outcomes, large crustacean specialists are the ancestral feeding regime, and all other feeding regimes evolve from this ancestral specialization. Compared to the rate correlation matrix estimated over the whole tree, large crustacean specialists present strong negative correlations between haploneme elongation and heteroneme size, and between rhopaloneme elongation and tentillum size, as well as with involucrum length. Within generalist clades (*Forskalia* and the *Agalma-Athorybia* clade), terminal filament nematocyst (desmoneme and rhopaloneme) sizes became negatively correlated with the sizes of most characters, meaning that as some tentilla became larger, their individual terminal nematocysts became smaller, observed to the extreme in *Agalma*. In addition, heteroneme and rhopaloneme elongation became positively correlated with cnidoband size. When large crustacean specialists switched to small crustacean prey in *Cordagalma* and calycophorans, haploneme size became inversely correlated with heteroneme elongation, which in turn developed a strong positive relationship with tentillum size. The extremes of this gradient can be seen in *Cordagalma* and *Hippopodius*, genera subspecialized in copepods and ostracods respectively. With the evolution of fish prey specialization in cystonects and within Clade B (Fig. 1), haploneme elongation became negatively correlated with heteroneme elongation (signal driven by Clade B, since cystonects lack tentacular heteronemes), and the surface area to volume ratio of haploneme nematocysts switched from a strong negative relationship with cnidoband size (found in every other regime) to a positive correlation. This is consistent with Clade B haplonemes becoming rounder, more similar to Cystonect haplonemes specialized in fish prey penetration and envenomation. Gelatinous specialization, albeit appearing only once in our tree, also carries a unique signature in character rate correlation shifts, with an increase in the strength of the correlation between heteroneme shape and shaft width, consistent with the appearance of birrhopaloid nematocysts with swollen shafts that are likely effective at anchoring gelatinous tissue (see reference to Narcomedusae nematocysts in (26)).

## Discussion

Several studies (12–16) have suggested that resource specialization can be an irreversible dead-end due to the constraints posed by extreme phenotypic specialization. Our results show that this is not the case for siphonophores, where the prey type on which they specialize has shifted at least 5 times. We find no support for any transitions from generalist to specialist (scenario 1, as described in the Introduction). We do find support for at least 3 instances of specialists switching from one prey type to another prey type, (scenario 2) and two switches from specialist to generalist (scenario 3). This is consistent with the findings of recent studies on phytophagous insects (19), where the rate of evolution from generalists to specialists is comparable to the reverse, thus specialization does not limit further evolution. Our results are also consistent with analyses of lepidopterans (21), where specialized resource switching is the primary transition type while niche breadth remains fairly constant. The evolutionary history of tentilla shows that siphonophores are an example of trophic niche diversification via morphological innovation and evolution, which allowed transitions between specialized trophic niches. In more familiar predators, the prey capture apparatus (such as claws and jaws) is well integrated in the body, leading to trade-offs and whole-body adaptations for feeding specialization. The extreme modularity of the siphonophore prey capture apparatus could release them from constraints typically imposed by adaptation to ecological specialization. This evolutionary mechanism is particularly important in a deep open ocean ecosystem, which is a relatively homogeneous physical environment, where the primary niche heterogeneity available is the potential interactions between organisms (8).

While selection acting on character states is a widely studied phenomenon, recent studies have shown that selection can also act upon the patterns of character correlations and phenotypic dependencies (33–39). This evolution of character relationships can allow lineages to explore new regions of the morphospace and facilitate the appearance of ecological novelties. Our results show that the patterns of phenotypic integration in siphonophore tentilla vary among clades, and appear to display different relationships across shifting feeding specializations. Similar to what has been found in the feeding morphologies of fish (33, 40), siphonophore tentilla may have accommodated new diets by altering the correlations between characters. For example, changes in the size and shape relationships between nematocyst types gave rise to the nematocyst complements specialized in ensnaring prey with different combinations of defensive traits.

Our results unambiguously show that tentillum morphology evolved with diet and strongly support deviations from the generalist-to-specialist evolution scenario. However, the conclusions we can draw from these analyses are limited in several ways. The biggest challenge at present is the sparse dietary data available in the literature. Additional dietary data could reveal transitions from generalists to specialists we were unable to detect for two reasons. First, some of the taxa in our dataset have a very limited number of feeding observations, which could lead to apparent specialization. Second, some of the taxa not included in our dataset could be undiscovered generalists. When interpreting these results, it is also important to remember that diet is also dependent on environmental prey availability. In addition, selectivity differences across siphonophore species could be also driven by other phenotypes not accounted for in this study. Finally, further observations on behavior, digestion biochemistry, and toxin composition are necessary to assess their relative importance in determining diet.

## Conclusions

Most studies on the evolution of predation have focused on vertebrate systems with an integrated feeding apparatus serving multiple functions. This has led to a narrow understanding of the evolutionary outcomes of specialization, where extreme morphological evolution constrains further shifts in their ecology. Siphonophores differ in many ways from commonly-known predators, using modular weapons for prey capture (the tentilla) that are fully decoupled from other structures and body functions. Our analysis of the evolutionary history of dietary specialization and morphological change in these elusive animals has revealed notable deviations from traditional expectations. While much of the feeding ecology literature focuses on how predatory generalists evolve into predatory specialists, in siphonophores we find predatory specialists can evolve into generalists, and that specialists on one prey type have directly evolved into specialists on other prey types. We find that the character states, evolutionary optima, and genetic correlations of many morphological characters have evolved following these ecological shifts. We find that the relationships between form and ecology hold across a large set of siphonophore taxa and characters. These findings are central to understanding the evolutionary mechanisms driving the emergence of food web complexity.

## Materials and Methods

### Tentillum morphology

The morphological work was carried out on siphonophore specimens fixed in 4% formalin from the Yale Peabody Museum Invertebrate Zoology (YPM-IZ) collection (accession numbers in Dryad repository). These specimens were collected intact across many years of fieldwork expeditions, using blue-water diving (41), remotely operated vehicles (ROVs), plankton net trawls, and human-operated submersibles. Tentacles were dissected from non-larval gastrozooids, sequentially dehydrated into 100% ethanol, cleared in methyl salicylate, and mounted onto slides with Canada Balsam or Permount mounting media. The slides were imaged as tiled z-stacks using differential interference contrast (DIC) on an automated stage at YPM-IZ (with the assistance of Daniel Drew and Eric Lazo-Wasem) and with laser point confocal microscopy using a 488 nm Argon laser that excited autofluorescence in the tissues. Thirty characters (defined in S1) were measured using Fiji (42, 43). We did not measure the lengths of contractile structures (terminal filaments, pedicles, gastrozooids, and tentacles) since they are too variable to quantify. We measured at least one specimen for 96 different species (raw data available in Dryad). Of these, we selected 38 focal species across clades based on specimen availability and phylogenetic representation. Three to five tentacle specimens from each one of these selected species were measured to capture intraspecific variation.

### Siphonophore phylogeny

While the main goal of this work is not to elucidate a novel phylogeny for Siphonophora, we did expand on the most recent transcriptome based phylogeny (24) to accommodate a larger taxon sampling. In order to do this, we ran a constrained analysis on an extensive 18S+16S dataset. The phylogenetic analysis included 55 siphonophore species and 6 outgroup cnidarian species (*Clytia hemisphaerica, Hydra circumcincta, Ectopleura dumortieri, Porpita porpita, Velella velella, Staurocladia wellingtoni*). The gene sequences we used in this study are available online (accession numbers in Dryad repository). Some of the sequences we used were accessioned in (27), and others we extracted from the transcriptomes in (24). Two new 16S sequences for *Frillagalma vityazi* (MK958598) and *Thermopalia* sp. (MK958599) sequenced by Lynne Christianson using the primers from (44) (read 3’ to 5’ F: TCGACTGTTTACCAAAAACATAGC, R: ACGGAATGAACTCAAATCATGTAAG) were included and accessioned to NCBI. Additional details on the phylogenetic inference methods can be found in the Supplementary Methods.

Unconstrained ML and Bayesian phylogenies were congruent (S2,S5). Given the broader sequence sampling of the transcriptome phylogeny, we ran constrained inferences (using both ML and Bayesian approaches, which produced fully congruent topologies (S4,S6)) after clamping the 5 nodes (S3) that were incongruent with the topology of the consensus tree in (24). This topology was then used to inform a Bayesian relaxed molecular clock time-tree in RevBayes, using a birth-death process (sampling probability calculated from the known number of described siphonophore species) to generate ultrametric branch lengths (S7-8). Scripts and tree files available in the Dryad repository.

### Feeding ecology

We extracted categorical diet data for different siphonophore species from published sources, including seminal papers (4, 25, 32, 45–48), and ROV observation data (22, 49) with the assistance of Elizabeth Hetherington and C. Anela Choy (data available in Dryad repository). In order to detect coarse-level patterns in feeding habits, the data were merged into feeding guilds. For more details on how the diet data was curated and summarized into guilds, please see Supplementary Methods.

We also extracted copepod prey length data from (25). To calculate specific prey selectivities, we extracted quantitative diet and zooplankton composition data from (32), matched each diet assessment to each prey field quantification by site, calculated Ivlev’s electivity indices (50), and averaged those by species (data available in Dryad repository).

### Statistical analyses

We used a series of phylogenetic comparative methods to test the evolutionary hypotheses presented in this study. We reconstructed ancestral states using ML (R phytools::anc.ML (51)), and stochastic character mapping (R phytools::make.simmap) for categorical characters. For more details on the data wrangling prior to these analyses, please see the Supplementary Methods. R scripts available in the Dryad repository.

In order to study the evolution of predatory specialization, we reconstructed components of the diet and prey selectivity on the phylogeny using ML (R phytools::anc.ML). To identify evolutionary associations of diet with tentillum and nematocyst characters, we compared the performance of a neutral evolution model to that of a diet-driven directional selection model. First, we collapsed the diet data into the five feeding guilds mentioned above (fish specialist, small crustacean specialist, large crustacean specialist, gelatinous specialist, generalist), based on which prey types they were observed consuming most frequently. Then, we reconstructed the feeding guild ancestral states using the ML function ace (package ape (52)), removing tips with no feeding data. The ML reconstruction was congruent with the consensus stochastic character mapping (S15). Then, using the package *OUwie* (53), we fitted an OU model with multiple optima and rates of evolution (OUm) matched to the reconstructed ancestral diet regimes, a single optimum OU model, and a BM null model, inspired by the analyses in (54). We then ranked the models in order of increasing parametric complexity (BM, OU, OUm), and compared the corrected Akaike Information Criterion (AICc) support scores (55) to the lowest (best) score, using a cutoff of 2 units to determine significantly better support. When the best fitting model was not significantly better than a less complex alternative, we selected the least complex model (S9). In addition, we calculated and reported the model adequacy scores using the R package *arbutus* (56).

In order to study correlations between the rates of evolution between different characters, we fitted a set of evolutionary variance-covariance matrices (33) (R phytools::evol.vcv). For more details on the data wrangling preceding these analyses, please see Supplementary Methods. To test whether phenotypic integration changed across selective regimes determined by the reconstructed feeding guilds, we carried out character-pairwise variance-covariance analysis comparing alternative models (R phytools::evolvcv.lite), including those where correlations are the same across the whole tree and models where correlations differ between selective regimes (S19). Number of taxa used in each pairwise comparison is reported in S20. Finally, we compared regime-specific variance-covariance matrices to the general matrix and to their preceding regime matrix to identify the changes in character dependences unique to each regime (S21-22).

We carried out a linear discriminant analysis of principal components (DAPC) using the dapc function (R adegenet::dapc) (57). This function allowed us to incorporate more predictors than individuals. We generated discriminant functions for feeding guild, and for the presence of copepods, fish, and shrimp (large crustaceans) in the diet (S10-13). From these DAPCs we obtained the highest contributing morphological characters to the discrimination (characters in the top quartile of the weighted sum of the linear discriminant loadings controlling for the eigenvalue of each discriminant). In order to identify the sign of the relationship between the predictor characters and prey type presence in the diet, we then generated generalized logistic regression models (as a type of generalized linear model, or GLM using R stats::glm) and phylogenetic generalized linear models (R phylolm::phyloglm) with the top contributing characters (from the corresponding DAPC) as predictors (S14). We also carried out these GLMs on the Ivlev’s selectivity indices for each prey type calculated from (32). In addition, we ran a series of comparative analyses to address hypotheses of diettentillum relationships posed in the literature. Additional details on the DAPC optimization are available in the Supplementary Materials.

## Supporting information

Collated PDF of supplementary methods and figures.

## Supplementary Materials

Data available from the Dryad Digital Repository: http://dx.doi.org/10.5061/dryad.NNNN Supplementary Materials are available in https://github.com/dunnlab/tentilla_morph/Supplement_forSupershort.pdf

## Acknowledgements

This work was supported by the Society of Systematic Biologists (Graduate Student Award to A.D.S.); the Yale Institute of Biospheric Studies (Doctoral Pilot Grant to A.D.S.); and the National Science Foundation (Waterman Award to C.W.D., and NSF-OCE 1829835 to C.W.D., S.H.D.H., and C. Anela Choy). A.D.S. was supported by a Fulbright Spain Graduate Studies Scholarship. We wish to thank the crew and scientists of the R/V Western Flyer, who participated in the collection of many of the specimens used in this study. We also want to thank Lynne Christianson and Shannon Johnson from the Monterey Bay Aquarium Research Institute for their assistance in the field as well as for sequencing some of the species included in this phylogeny. In addition, we wish to thank Lourdes Rojas, Daniel Drew, and Eric Lazo-Wasem for their assistance in imaging the fixed specimens and managing the collections. We thank Dennis Pilarczyk for organizing the prey selectivity data, Michael Landis for helping design the Bayesian analyses, and Joaquin Nunez for reviewing this manuscript. Furthermore, we thank Elizabeth D. Hetherington and C. Anela Choy for collating the data on siphonophore feeding and for reviewing the manuscript. Finally, we thank Philip Pugh, who confirmed many of our specimen identifications and taught us valuable knowledge about siphonophores.

