## Supplementary material for "Shaped to kill: The evolution of siphonophore tentilla for specialized prey capture in the open ocean": Collated PDF of supplementary methods and figures.

### Supplementary Materials File

#### Supplementary Methods

Phylogenetic inference: We aligned the sequences using MAFFT (1) (alignments available in Dryad). We inferred a Maximum Likelihood (ML) phylogeny (S2) from 16S and 18S ribosomal rRNA genes using IQTree (2) with 1000 bootstrap replicates (iqtree -s alignment.fa -nt AUTO -bb 1000). We used ModelFinder (3) implemented in IQTree v1.5.5. to assess the relative model fit. ModelFinder selected GTR+R4 for having the lowest Bayesian Information Criterion score. Additionally, we inferred a Bayesian tree with each gene as an independent partition in RevBayes (4) (S5), which was topologically congruent with the unconstrained ML tree. The *alpha* priors were selected to minimize prior load in site variation.

Diet data curation: We removed the gelatinous prey observations for *Praya dubia* eating a ctenophore and a hydromedusa, and for *Nanomia* sp. eating *Aegina* since we believe these are rare events that have a much larger probability of being detected by ROV methods than their usual prey, and it is not clear whether the medusae were attempting to prey upon the siphonophores. Personal observations on feeding (from SHDH, Anela Choy, and Philip Pugh) were also included for *Resomia ornicephala*, *Lychnagalma utricularia*, *Bargmannia amoena*, *Erenna richardi*, *Erenna laciniata*, *Erenna sirena*, and *Apolemia rubriversa*. The feeding guilds declared in this study are: small-crustacean specialist (feeding mainly on copepods and ostracods), large crustacean specialist (feeding on large decapods, mysids, or krill), fish specialist (feeding mainly on actinopterygian larvae, juveniles, or adults), gelatinous specialist (feeding mainly on other siphonophores, medusae, ctenophores, salps, and/or doliolids), and generalist (feeding on a combination of the aforementioned taxa, without favoring any one prey group). These were selected to minimize the number of categories while keeping the most different types of prey separate.

Data wrangling for comparative analyses: For comparative analyses, we removed species present in the tree but not represented in the morphology data, and *vice versa*. Although we measured specimens labeled as *Nanomia bijuga* and *Nanomia cara*, we are not confident in some of the species-level identifications, and some specimens were missing diagnostic zooids. Thus, we decided to collapse these into a single taxonomic concept (*Nanomia* sp.). All *Nanomia* sp. observations were matched to the phylogenetic position of *Nanomia bijuga* in the tree. We carried out all phylogenetic comparative statistical analyses in the programming environment R (5), using the Bayesian ultrametric species tree (S8), and incorporating intraspecific variation estimated from the specimen data as standard error whenever the analysis tool allowed it. R scripts and summarized species-collapsed data available in the Dryad repository. For each character (or character pair) analyzed, we removed species with missing data and reported the number of taxa included. We tested each character for normality using the Shapiro-Wilk test (6), and log-transformed those that were non-normal.

Data wrangling for the variance-covariance analyses: When fitting all variance-covariance terms simultaneously (S16-18), we selected the largest set of characters that would allow the analysis to run without computational singularities. This excluded many of the morphometric characters which are linearly dependent on other characters. Since the functions do not tolerate missing data, we ran the analyses in two ways: One including all taxa but transforming absent states to zeroes, and another removing the taxa with absent states. These analyses could only be carried out on the subset of taxa for which diet data is available, and only among character pairs that are not computationally singular for that taxonomic subset. Gelatinous specialist correlations could only be estimated for a small subset of characters present in *Apolemia* (S21F, S22E, S23D) and should be interpreted with care.

Comparative tools used to test character associations: To test for correlated evolution among binary characters, we used Pagel's test (7). To characterize and evaluate the relationship between continuous characters, we used phylogenetic generalized least squares regressions (PGLS) (8). To compare the evolution of continuous characters with categorical aspects of the diet, we carried out a phylogenetic logistic regression (R nlme::gls using the 'corBrownian' function for the argument 'correlation').

DAPC optimization: Some taxa have inapplicable states for certain absent characters (such as the length of a nematocyst subtype that is not present in a species), which are problematic for DAPC analyses. We tackled this by transforming the absent states to zeroes. This approach allows us to incorporate all the data, but creates an attraction bias between small character states (*e.g.* small tentilla) and absent states (*e.g.* no tentilla). Absent characters are likely to be very biologically relevant to prey capture and we believe they should be accounted for in a predictive approach. We limited the number of linear discriminant functions retained to the number of groupings in each case. We selected the number of principal components retained using the a-score optimization function (R adegenet::optim.a.score) (9) with 100 iterations, which yielded more stable results than the cross validation function (R adegenet::xval). This optimization aims to find the compromise value with highest discrimination power with the least overfitting. The discriminant analysis for feeding guild (7 principal components, 4 discriminants) produced 100% discrimination, and the highest loading contributions were found for the characters (ordered from highest to lowest): Involucrum length, heteroneme volume, heteroneme number, total heteroneme volume, tentacle width, heteroneme length, total nematocyst volume, and heteroneme width (S10).

### Supplementary Materials

**2.1) Definitions of homologous structures used throughout this work.**

| Structure | Definition |
|---|---|
| **Haploneme** | Nematocyst with no shaft |
| **Heteroneme** | Nematocyst with a distinct shaft |
| **Desmoneme** | Small oval/tapered adhesive nematocyst with thick coiled tubule |
| **Rhopaloneme** | Small rod-like nematocyst found on the terminal filament |
| **Terminal filament** | Distal extension of the tentillum beyond the cnidoband |
| **Cnidoband** | Distinct packing of nematocysts on the dorsal side of the tentillum |
| **Tentacle** | Tubular projection from the gastrozooid basigaster |
| **Tentillum** | Evenly spaced dorsal evagination of the tentacle carrying ordered and functional nematocysts |
| **Involucrum** | Extension of the pedicle covering part of the cnidoband |
| **Pedicle** | Proximal region of the tentillum between the cnidoband and the tentacle |
| **Elastic strand** | Mesoglea derived collagenous double strand underlying the cnidoband of some siphonophores |

**2.2) Definitions of the continuous morphological and kinematic characters measured.**

| Character | Definition | Units |
|---|---|---|
| **Cnidoband length** | Distance from the base to the tip of the cnidoband in natural position | micrometers |
| **Cnidoband free length** | Distance from the base to the tip of the cnidoband when stretched straight | micrometers |
| **Cnidoband width** | Diameter of the cnidoband on the widest point | micrometers |
| **Involucrum length** | Length of the involucrum from the base of the cnidoband to its most distal extent | micrometers |
| **Heteroneme length** | Length of the heteronemes | micrometers |
| **Heteroneme width** | Diameter of the heteronemes at the widest point | micrometers |
| **Heteroneme shaft length** | Length of the heteroneme shaft | micrometers |
| **Heteroneme shaft width** | Width of the heteroneme shaft | micrometers |
| **Heteroneme number** | Number of heteronemes in each tentillum (# in each row*2) | micrometers |
| **Haploneme length** | Length of the haplonemes | micrometers |
| **Haploneme width** | Diameter of the haplonemes at the widest point | micrometers |
| **Rhopaloneme length** | Length of the rhopalonemes | micrometers |
| **Rhopaloneme width** | Diameter of the rhopalonemes at the widest point | micrometers |
| **Desmoneme length** | Length of the desmonemes | micrometers |
| **Desmoneme width** | Diameter of the cnidoband at the widest point | micrometers |
| **Involucrum length** | Length of the involucrum from the base of the cnidoband to its most distal extent | micrometers |
| **Elastic strand width** | Diameter of the descending elastic strand at the widest point | micrometers |
| **Pedicle width** | Diameter of the pedicle | micrometers |
| **Tentacle width** | Diameter of the tentacle | micrometers |
| **Haploneme row number** | Number of haploneme rows running parallel to the length of the cnidoband | micrometers |
| **Cnidoband coiledness** | Cnidoband free length / Cnidoband length | adimensional |
| **Heteroneme elongation** | Heteroneme Length/Width | adimensional |
| **Haploneme elongation** | Haploneme Length/Width | adimensional |
| **Desmoneme elongation** | Desmoneme Length/Width | adimensional |
| **Rhopaloneme elongation** | Rhopaloneme Length/Width | adimensional |
| **Heteroneme shaft extension** | Heteroneme shaft length / Heteroneme capsule length | adimensional |
| **Nematocyst Surface area** | 4*pi*(2*((((Length/2)*(Width/2))^1.6)+(((Width/2)^2)^1.6))/3)^(1/1.6) | micrometers squared |
| **Nematocyst volume** | Ellipsoid formula : (4/3)*pi*(Length/2)*((Width/2)^2) | micrometers cubed |
| **Nematocyst SA/V ratio** | Nematocyst surface area / Nematocyst volume | 1/micrometers |
| **Total haploneme volume** | Haploneme volume * Haploneme row number * (Cnidoband free length / Haploneme width) | micrometers cubed |
| **Total heteroneme volume** | Heteroneme volume * Heteroneme number | micrometers cubed |
| **Total nematocyst volume** | Total haploneme volume + Total heteroneme volume | micrometers cubed |
| **Total discharge time** | Time from initial cnidoband movement to complete conformational change | miliseconds |
| **Average CB discharge speed** | Distance covered by the leading edge of the discharging cnidoband in the total discharge time. | mm/s |
| **Maximum CB discharge speed** | Maximum speed attained by the leading edge of the discharging cnidoband | mm/s |
| **Heteroneme discharge speed AVG** | Distance covered by the heteroneme nematocyst tubule from initial ejection to full eversion in the time it takes to evert fully. | mm/s |
| **Heteroneme discharge free speed AVG** | Distance covered by the heteroneme nematocyst tubule in the time it takes to evert fully, accounting for coiling. | mm/s |
| **Heteroneme discharge speed MAX** | Maximum speed attained by the non-shaft tubule of the heteroneme nematocysts during eversion. | mm/s |
| **Heteroneme discharge free speed MAX** | Maximum speed attained by the non-shaft tubule of the heteroneme nematocysts during eversion, accounting for coiling. | mm/s |
| **Heteroneme shaft discharge speed MAX** | Maximum speed attained by the shaft of the tubule of the heteroneme nematocysts during initial eversion. | mm/s |
| **Heteroneme filament length** | Distance covered by the heteroneme nematocyst tubule from initial ejection to full eversion | micrometers |
| **Haploneme discharge speed AVG** | Distance covered by the haploneme nematocyst tubule from initial ejection to full eversion in the time it takes to evert fully. | mm/s |

Figure 1: Character definitions.

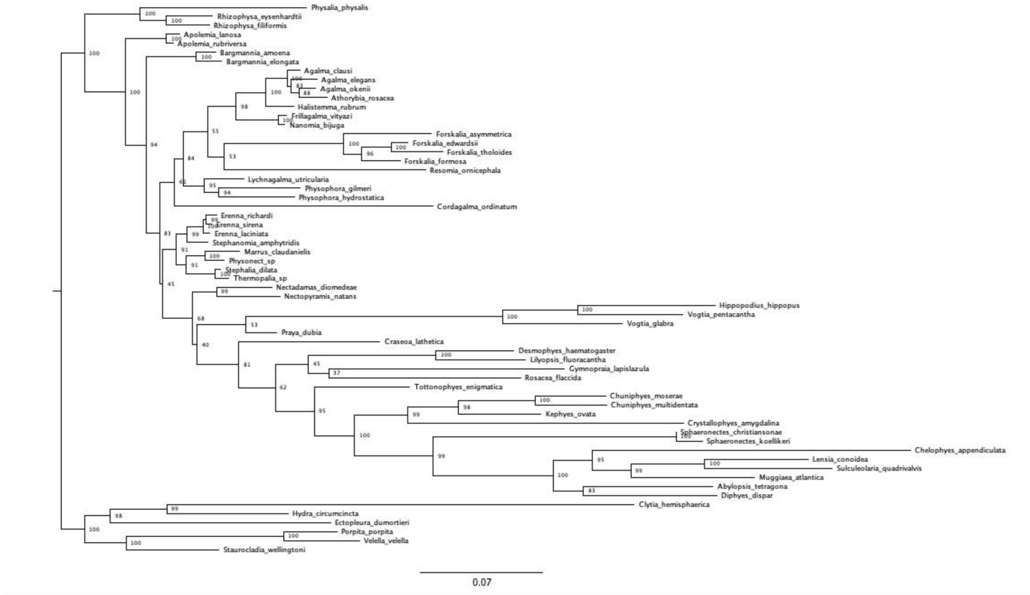


Figure 2: Maximum likelihood IQTree inference, unconstrained. Node labels are bootstrap support values.

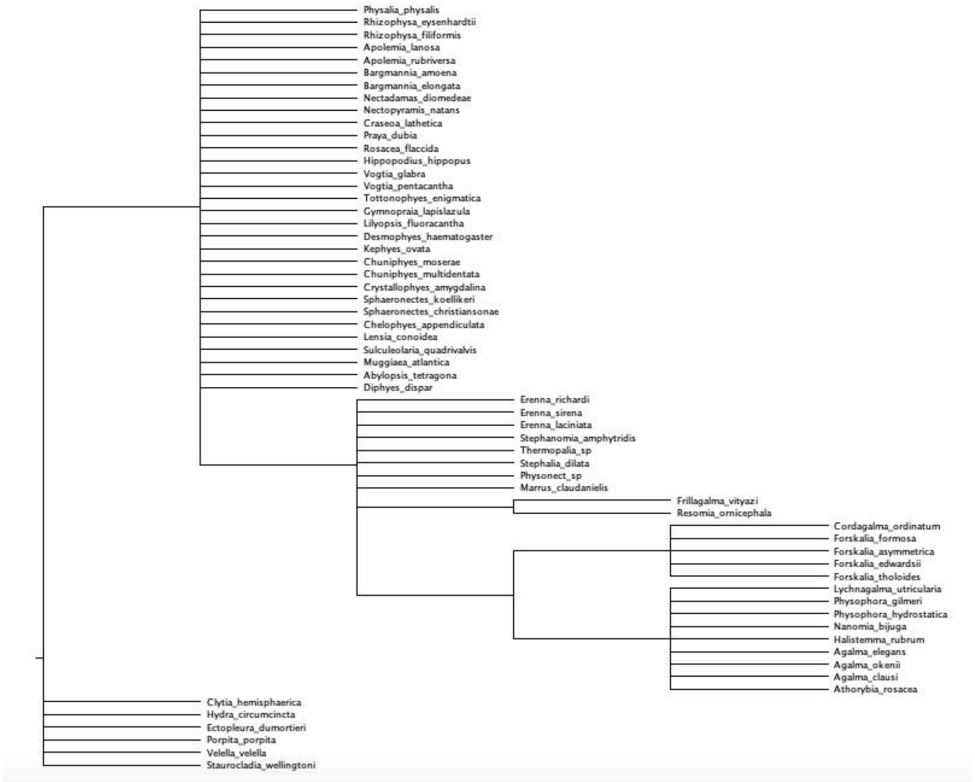


Figure 3: Topology used to constrain analyses (minimal topological statements based on the incongruences between the unconstrained tree and Munro et al. (2018).

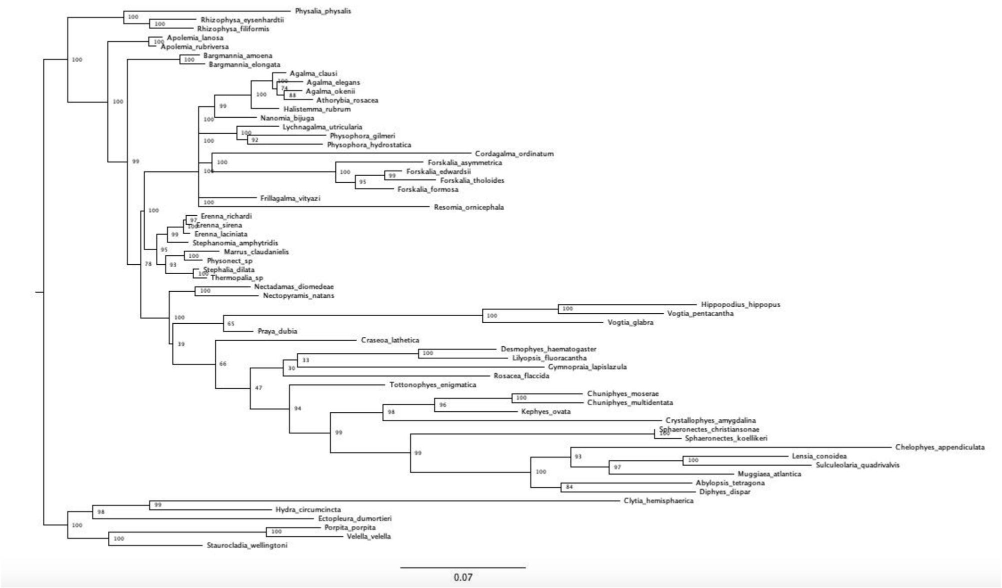


Figure 4: Constrained IQTree ML inference. Node labels are bootstrap support values.

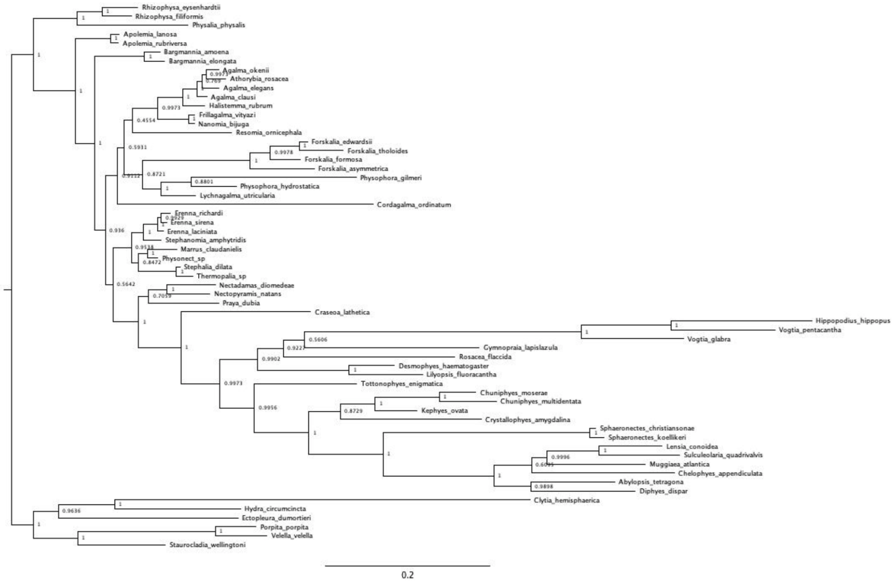


Figure 5: Unconstrained Bayesian topology inference in RevBayes (node labels are Bayesian posteriors).

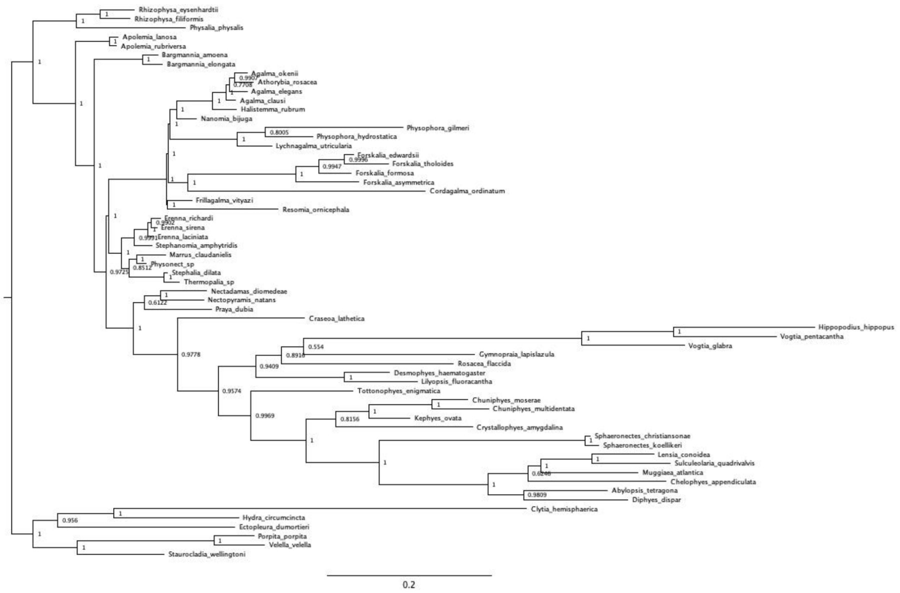


Figure 6: Clade constrained Bayesian inference in RevBayes (node labels are Bayesian posteriors).

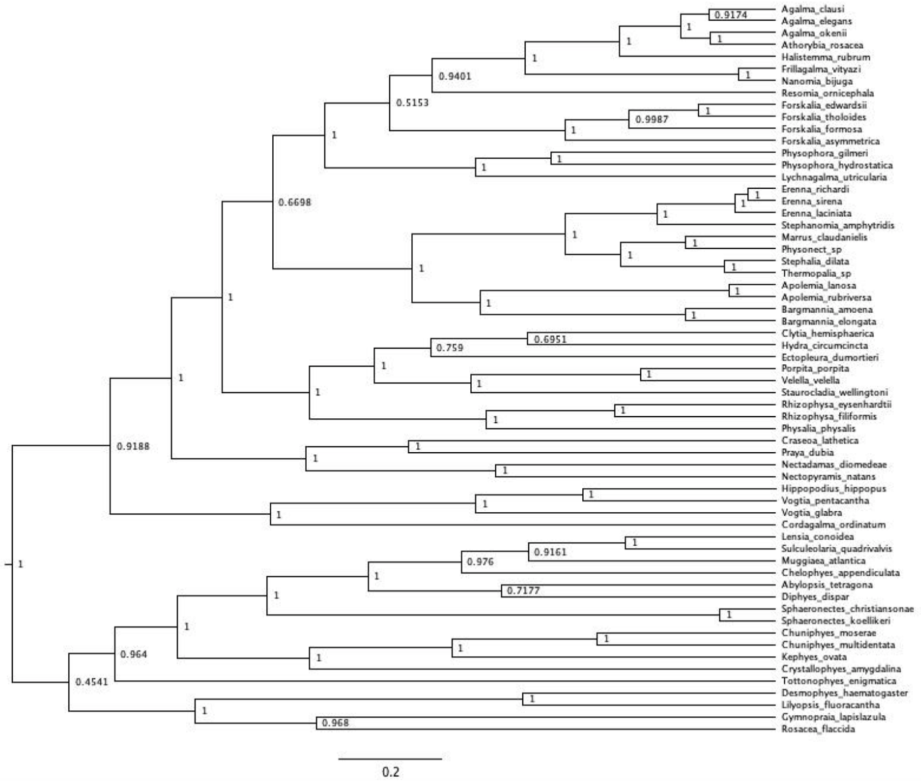


Figure 7: Unconstrained ultrametric Bayesian time tree branch length and topology inference in RevBayes (node labels are Bayesian posteriors). Arbitrary rooting.).

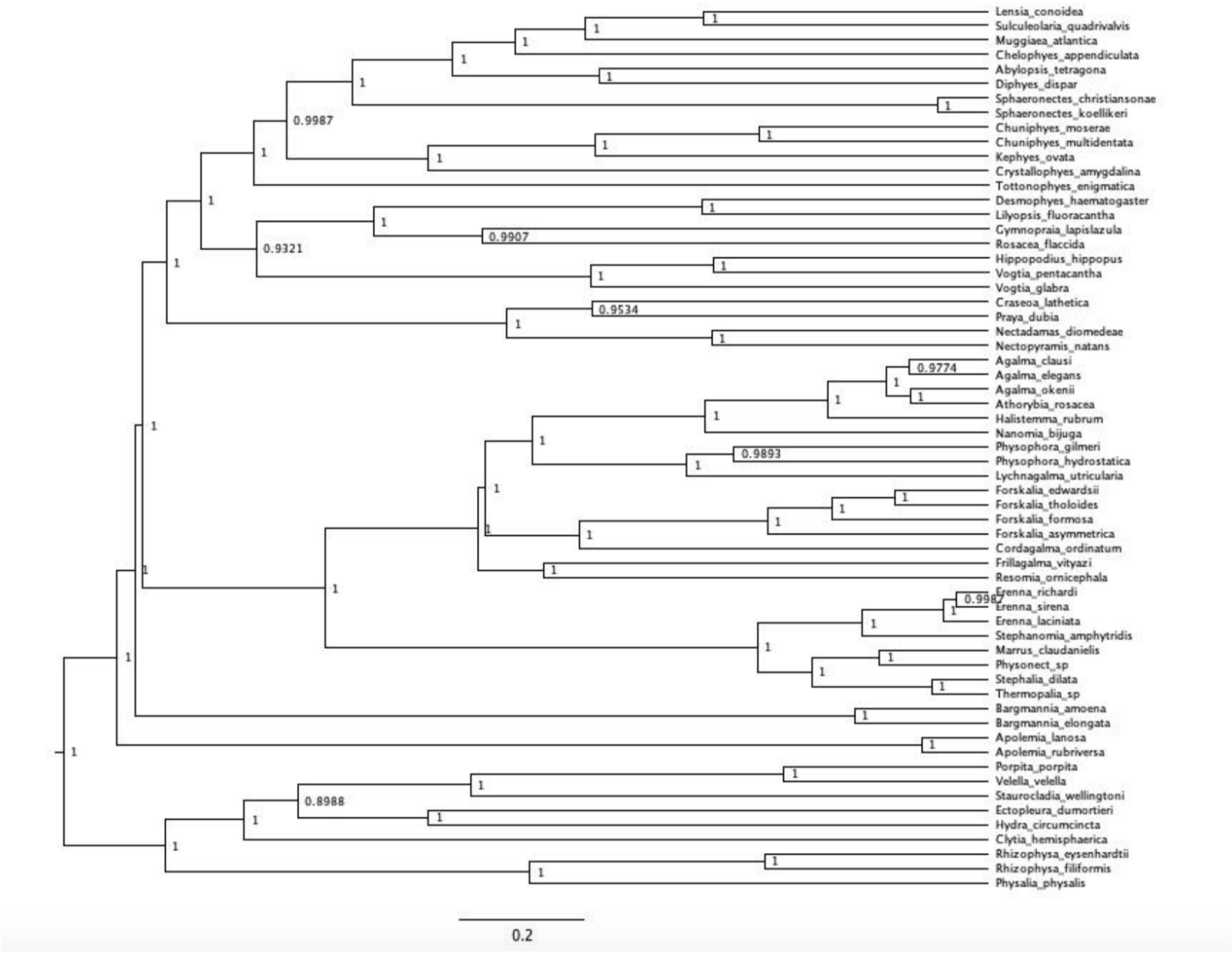


Figure 8: Ultrametric Bayesian time tree branch length inference in RevBayes (node labels are bayesian posteriors). Topology clamped to the Bayesian constrained topology inference in @ref{Bayes_constrained}. Tree rooted using outgroup constraint.

| Character | N | dAICc BM | dAICc OU1 | dAICc OUm | Msig | Cvar | Svar | Sasr | Shgt | Dcfd |
|---|---|---|---|---|---|---|---|---|---|---|
| Haploneme elongation | 21 | 0 | 0.953 | 713.671 | 0.801 | 0 | 0.038 | 0.156 | 0.362 | 0.098 |
| Heteroneme shaft width µm | 19 | 0 | 1.051 | 632.503 | 0.767 | 0.801 | 0.128 | 0.092 | 0.4 | 0.813 |
| Cnidoband width µm | 21 | 0 | 1.595 | 761.241 | 0.781 | 0.723 | 0.072 | 0.09 | 0.31 | 0.228 |
| Heteroneme shaft free length µm | 19 | 0 | 1.649 | 628.334 | 0.791 | 0.402 | 0.941 | 0.098 | 0.575 | 0.464 |
| Heteroneme volume µm3 | 19 | 0 | 2.105 | 629.21 | 0.779 | 0.034 | 0.39 | 0.338 | 0.637 | 0.392 |
| Haploneme width µm | 21 | 0 | 2.452 | 766.546 | 0.779 | 0.599 | 0.316 | 0.791 | 0.995 | 0.288 |
| Pedicle width µm | 21 | 0 | 2.458 | 764.406 | 0.815 | 0.791 | 0.368 | 0.26 | 0.963 | 0.298 |
| Heteroneme width µm | 19 | 0 | 2.516 | 634.229 | 0.805 | 0.809 | 0.292 | 0.208 | 0.709 | 0.38 |
| Tentacle width µm | 22 | 0 | 2.702 | 383.12 | 0.835 | 0.496 | 0.344 | 0.867 | 0.096 | 0.444 |
| Heteroneme to CB | 19 | 0 | 0.127 | NA | 0.811 | 0.336 | 0.004 | 0.068 | 0.026 | 0.434 |
| Haploneme surface area:volume | 21 | 0 | 2.282 | 757.267 | 0.747 | 0.563 | 0.392 | 0.583 | 0.927 | 0.15 |
| Heteroneme elongation | 19 | 0.217 | 0 | 618.621 | 0.819 | 0.601 | 0.012 | 0.707 | 0.062 | 0.715 |
| Total nematocyst volume | 22 | 0.57 | 0 | 378.872 | 0.809 | 0.501 | 0.06 | 0.088 | 0.266 | 0.501 |
| Heteroneme free length µm | 19 | 0.746 | 0 | 627.372 | 0.811 | 0.885 | 0.593 | 0.156 | 0.368 | 0.679 |
| Total haploneme volume | 21 | 1.281 | 0 | 730.592 | 0.829 | 0.452 | 0.038 | 0.134 | 0.096 | 0.819 |
| Cnidoband length µm | 21 | 1.439 | 0 | 763.478 | 0.761 | 0.328 | 0.04 | 0.11 | 0.098 | 0.803 |
| Cnidoband free length µm | 21 | 2.219 | 0 | 760.518 | 0.843 | 0.35 | 0.012 | 0.066 | 0.05 | 0.911 |
| Cnidoband coiledness | 21 | 2.669 | 0 | 765.921 | 0.807 | 0.002 | 0.008 | 0.03 | 0.076 | 0.791 |
| Haploneme row number | 21 | 4.177 | 0 | 729.95 | 0.825 | 0.004 | 0.002 | 0.06 | 0.006 | 0.346 |
| Haploneme free length µm | 21 | 5.497 | 0 | 778.011 | 0.793 | 0.388 | 0.032 | 0 | 0.052 | 0.306 |
| Heteroneme shaft extension | 19 | 6.17 | 0 | 611.533 | 0.775 | 0 | 0.068 | 0.665 | 0.124 | 0.184 |
| Rhopaloneme elongation | 13 | 144.229 | 146.783 | 0 | 0.753 | 0.641 | 0.434 | 0.188 | 0.933 | 0.617 |
| Desmoneme length µm | 13 | 148.14 | 151.403 | 0 | 0.763 | 0.182 | 0.607 | 0.31 | 0.745 | 0.014 |
| Rhopaloneme length µm | 13 | 150.731 | 154.198 | 0 | 0.739 | 0.803 | 0.24 | 0.03 | 0.14 | 0.316 |
| Rhopaloneme width µm | 13 | 150.82 | 154.287 | 0 | 0.743 | 0.462 | 0.306 | 0.07 | 0.182 | 0.092 |
| Desmoneme elongation | 13 | 159.594 | 158.584 | 0 | 0.719 | 0.206 | 0.074 | 0.094 | 0.036 | 0.993 |
| Desmoneme width µm | 13 | 164.639 | 168.106 | 0 | 0.773 | 0.11 | 0.885 | 0.098 | 0.605 | 0.002 |
| Involucrum length µm | 14 | 148.672 | 151.078 | 0 | 0.779 | 0.126 | 0.17 | 0.25 | 0.418 | 0.671 |
| Elastic strand width µm | 15 | 473.984 | 477.156 | 0 | 0.827 | 0.921 | 0.184 | 0.064 | 0.953 | 0.785 |
| Total heteroneme volume | 17 | 619.03 | 619.932 | 0 | 0.797 | 0.803 | 0.078 | 0.172 | 0.35 | 0.697 |
| Heteroneme number | 17 | 620.836 | 620.193 | 0 | 0.777 | 0.39 | 0.008 | 0.074 | 0.056 | 0.054 |

Brownian Motion Supported (rows Haploneme elongation – Cnidoband length µm)

Single Optimum OU Supported (rows Cnidoband free length µm – Heteroneme shaft extension)

Multiple Optima OU Supported (rows Rhopaloneme elongation – Heteroneme number)

Figure 9: Model support (delta AICc) for each morphological character analyzed on the feeding guild reconstruction regime tree. OU1 = Single-optimum Ornstein-Uhlenbeck. OUm = Multi-optima Ornstein-Uhlenbeck. Model adequacy scores calculated for the best supported model only. Msig = mean of squared contrasts. Cvar = coefficient of variation of the absolute value of the contrasts. Svar = Slope of a linear model fitted to the absolute value of the contrasts against their expected variances. Sasr = slope of the contrasts against the ancestral state inferred at each corresponding node. Shgt = slope of the contrasts against node depth. Dcfd = Kolmolgorov-Smirnov D-statistic comparing contrasts to a normal distribution with SD equal to the root of the mean of squared contrasts.

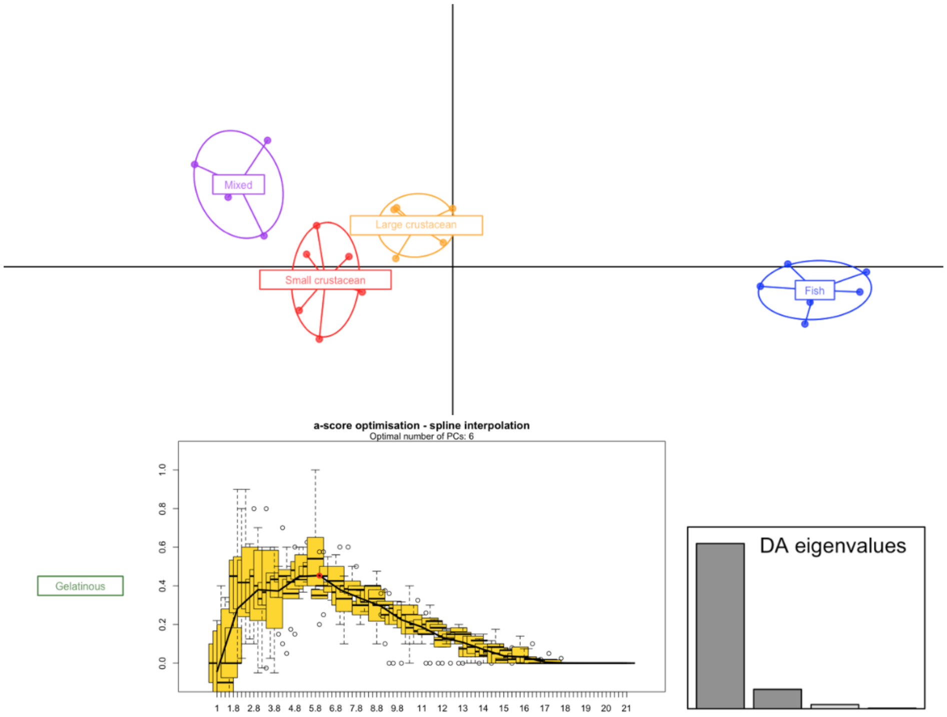


| Variable contribution | |
|---|---|
| Involucrum.length..um. | 91.945027 |
| Heteroneme.volume..um3. | 71.675594 |
| Heteroneme.number | 33.874805 |
| Total_heteroneme_volume | 17.913878 |
| Tentacle.width..um. | 13.274347 |
| Heteroneme.free.length..um. | 10.825676 |
| Total nematocyst volume | 9.838328 |
| Heteroneme.width..um. | 8.474876 |

Figure 10: DAPC for Feeding guilds. Six PCs retained after a-score optimization (100 iterations). Four LDA functions used. Discriminant power on training set: 100%. Prediction posterior distribution heat map in main text Figure 6. Variable contribution (top quartile) calculated by the sum of the LDA variable loadings weighted by the eigenvalue of each LDA.

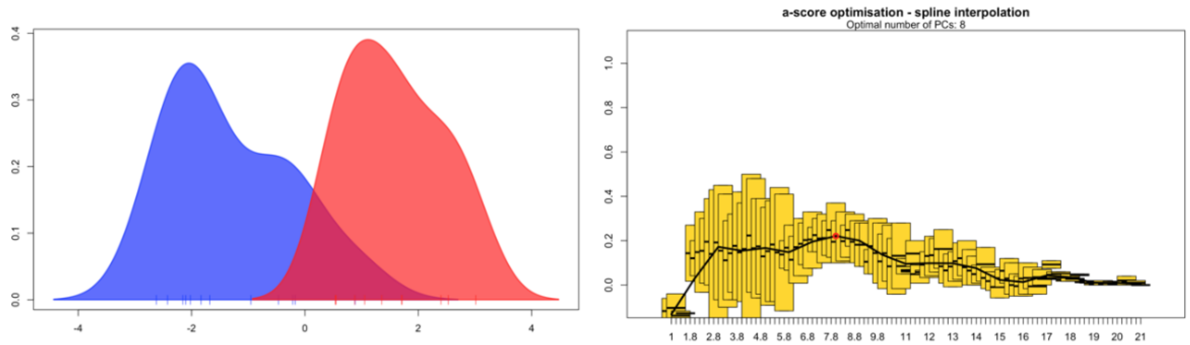


**Variable contribution**

| | |
|---|---|
| Total_nematocyst_volume | 12.810953 |
| Tentacle.width..um. | 5.687086 |
| haploneme_elongation | 4.586386 |
| SAV_haploneme | 4.264843 |
| Haploneme.row.number..um. | 2.966009 |
| Cnidoband.length..um. | 1.959479 |
| Cnidoband.width..um. | 1.679753 |
| Cnidoband.free.length..um. | 1.468262 |

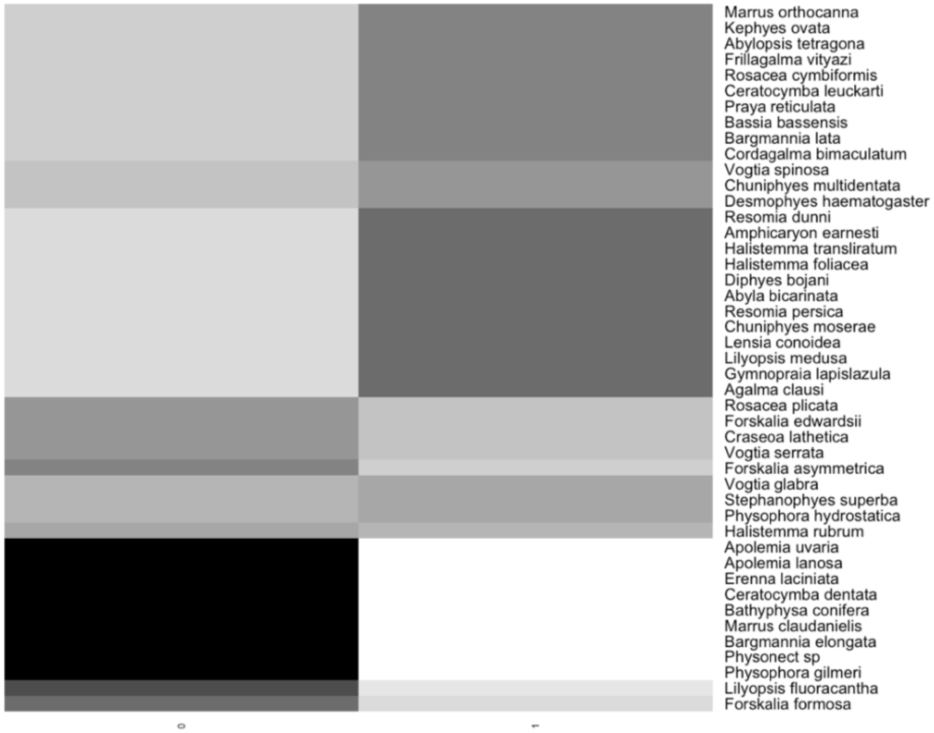


Figure 11: DAPC for copepod presence in the diet. Eight PCs retained after a-score optimization (100 iterations). One LDA functions used. Discriminant power on training set: 95.4%. Grayscale heat map shows the posterior probability distribution of the predictions. Variable contribution (top quartile) calculated by the sum of the LDA variable loadings weighted by the eigenvalue of each LDA.

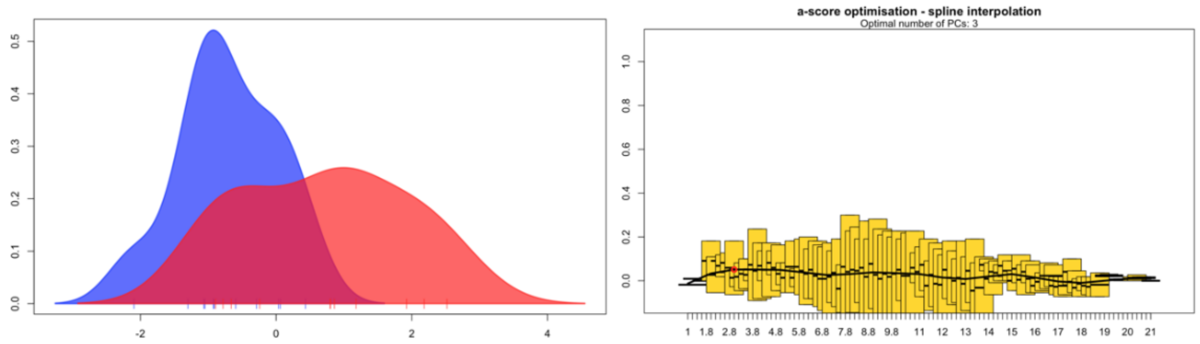


| | Variable contribution |
|---|---|
| total_haploneme_volume | 2.2734508 |
| Heteroneme.volume..um3. | 1.1308252 |
| total_nematocyst_volume | 1.1104459 |
| total_heteroneme_volume | 0.9402038 |
| Cnidoband.length..um. | 0.7583124 |
| Cnidoband.free.length..um. | 0.6650068 |
| Involucrum.length..um. | 0.6097537 |
| Pedicle.width..um. | 0.5447312 |

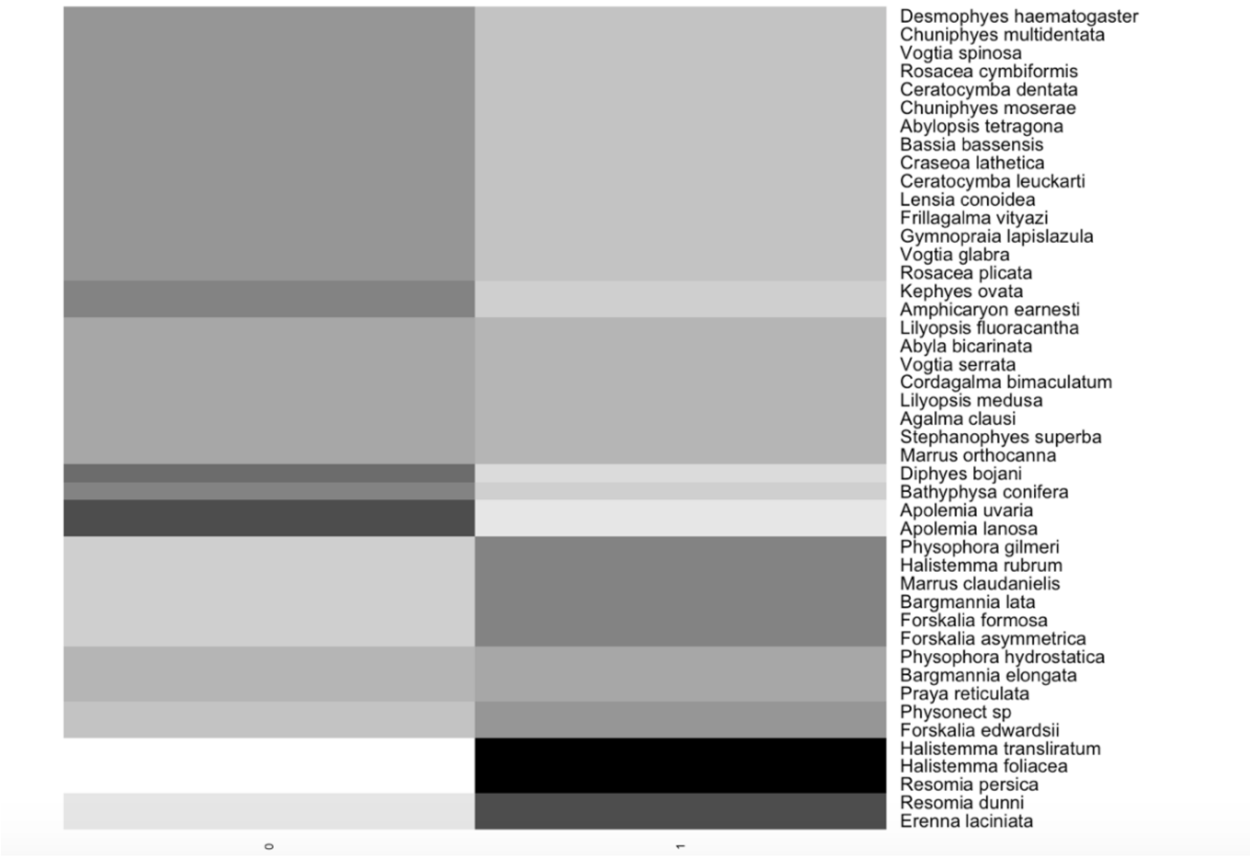


Figure 12: DAPC for fish presence in the diet. Three PCs retained after a-score optimization (100 iterations). One LDA function used. Discriminant power on training set: 68.1%. Grayscale heat map shows the posterior probability distribution of the predictions. Variable contribution (top quartile) calculated by the sum of the LDA variable loadings weighted by the eigenvalue of each LDA.

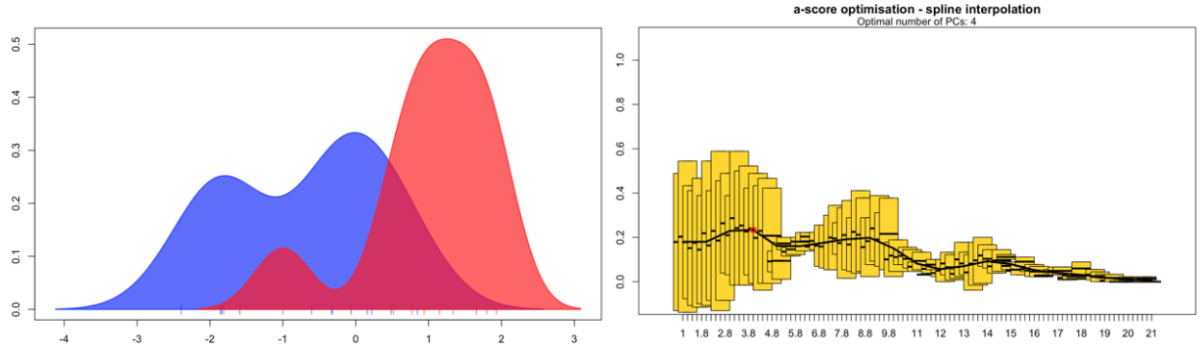


| | Variable contribution |
|---|---|
| Involucrum.length..um. | 8.4739326 |
| total_heteroneme_volume | 2.0479062 |
| Elastic.strand.width..um. | 1.2640038 |
| Rhopaloneme.length..um. | 0.4274179 |
| Heteroneme.volume..um3. | 0.4255758 |
| haploneme_elongation | 0.3530771 |
| Desmoneme.length..um. | 0.3274451 |
| Tentacle.width..um. | 0.2763979 |

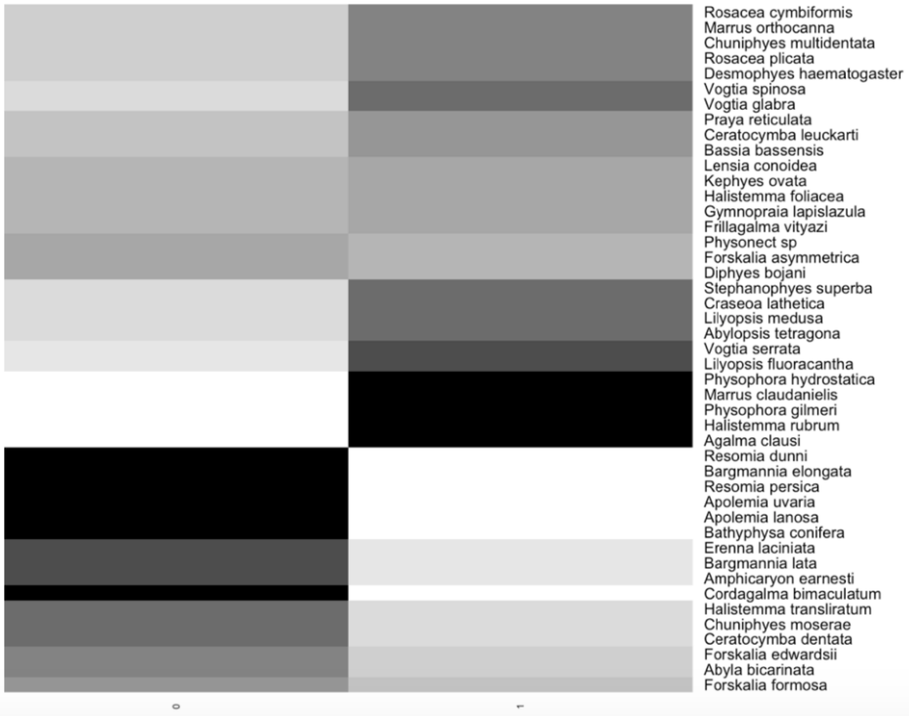


Figure 13: DAPC for large crustacean presence in the diet. Four PCs retained after a-score optimization (100 iterations). One LDA function used. Discriminant power on training set: 81.8%. Grayscale heat map shows the posterior probability distribution of the predictions. Variable contribution (top quartile) calculated by the sum of the LDA variable loadings weighted by the eigenvalue of each LDA.

| Character | Prey type | Ntaxa | phyloGLM AIC | phyloGLM P | phyloglm b | GLM AIC | GLM P | GLM b |
|---|---|---|---|---|---|---|---|---|
| **Cnidoband coiledness** | **Decapod diet** | 21 | 23.701 | 0.029 | 2.327 | 21.762 | 0.016 | 3.227 |
| **Haploneme surface area:volume** | **Copepod diet** | 21 | 19.143 | 0.017 | 3.246 | 17.355 | 0.017 | 4.631 |
| **Haploneme width µm** | **Copepod diet** | 21 | 18.844 | 0.017 | -3.098 | 16.997 | 0.019 | -4.417 |
| **Pedicle width µm** | **Copepod diet** | 21 | 22.182 | 0.032 | -1.16 | 23.723 | 0.024 | -1.437 |
| **Tentacle width µm** | **Copepod diet** | 22 | 22.038 | 0.026 | -1.543 | 23.634 | 0.025 | -1.505 |
| **Cnidoband length µm** | **Copepod diet** | 21 | 23.431 | 0.042 | -0.864 | 24.178 | 0.025 | -1.131 |
| **Cnidoband width µm** | **Copepod diet** | 21 | 22.887 | 0.035 | -1.545 | 23.658 | 0.027 | -1.89 |
| **Heteroneme number** | **Copepod diet** | 17 | 20.52 | 0.059 | -0.718 | 19.615 | 0.03 | -0.973 |
| **Total haploneme volume** | **Copepod diet** | 21 | 23.507 | 0.03 | -0.581 | 25.232 | 0.031 | -0.578 |
| **Total heteroneme volume** | **Copepod diet** | 17 | 17.156 | 0.032 | -0.533 | 16.369 | 0.031 | -0.758 |
| **Pedicle width µm** | **Ostracod diet** | 21 | 17.523 | 0.041 | -1.43 | 15.165 | 0.035 | -1.97 |
| **Heteroneme shaft free length µm** | **Copepod diet** | 19 | 23.955 | 0.076 | -1.53 | 23.378 | 0.04 | -2.16 |
| **Haploneme width µm** | **Fish diet** | 21 | 28.118 | 0.091 | 1.268 | 27.551 | 0.043 | 1.642 |
| **Tentacle width µm** | **Fish diet** | 22 | 28.927 | 0.058 | 0.804 | 28.771 | 0.044 | 0.874 |
| **Haploneme surface area:volume** | **Fish diet** | 21 | 28.258 | 0.098 | -1.329 | 27.596 | 0.044 | -1.768 |
| **Total haploneme volume** | **Ostracod diet** | 21 | 20.028 | 0.043 | -0.619 | 17.733 | 0.046 | -0.681 |
| **Heteroneme volume µm3** | **Copepod diet** | 19 | 24.282 | 0.091 | -0.521 | 24.297 | 0.046 | -0.72 |
| **Pedicle width µm** | **Fish diet** | 21 | 28.21 | 0.074 | 0.815 | 27.839 | 0.049 | 0.918 |

Figure 14: Logistic regressions between continuous morphological characters and prey type presences. Ntaxa = number of taxa used in the analyses after removing taxa with missing diet data and inapplicable character states. phyloGLM = Phylogenetic generalized logistic regression model. GLM = Generalized logistic regression model. P = p-value. b = slope. Only cases with significant GLM fits were retained. Cells colored blue indicate phyloGLM p-value $< 0.05$. Cells colored green indicate GLM p-value $< 0.05$

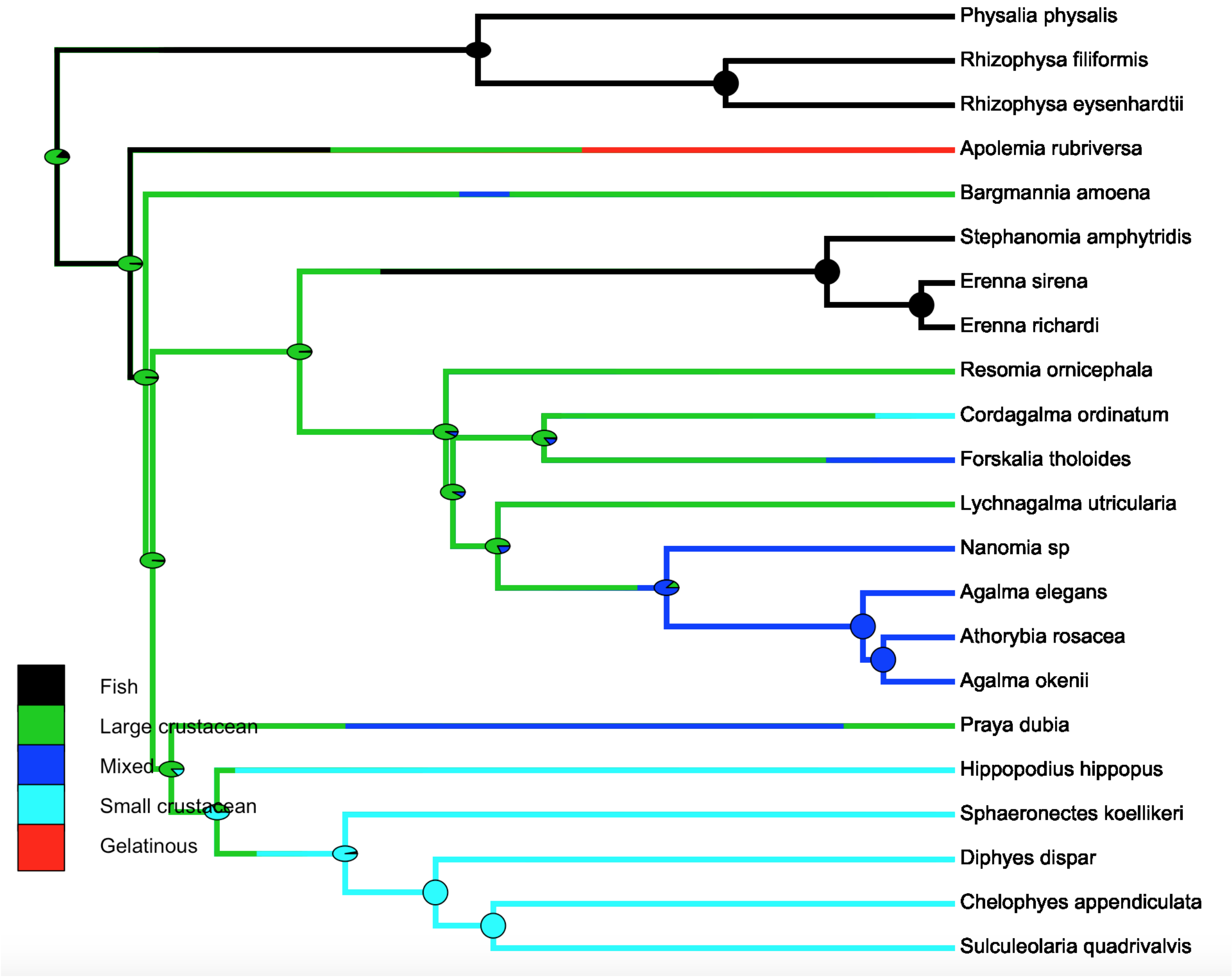


Figure 15: SIMMAP Feeding guilds.

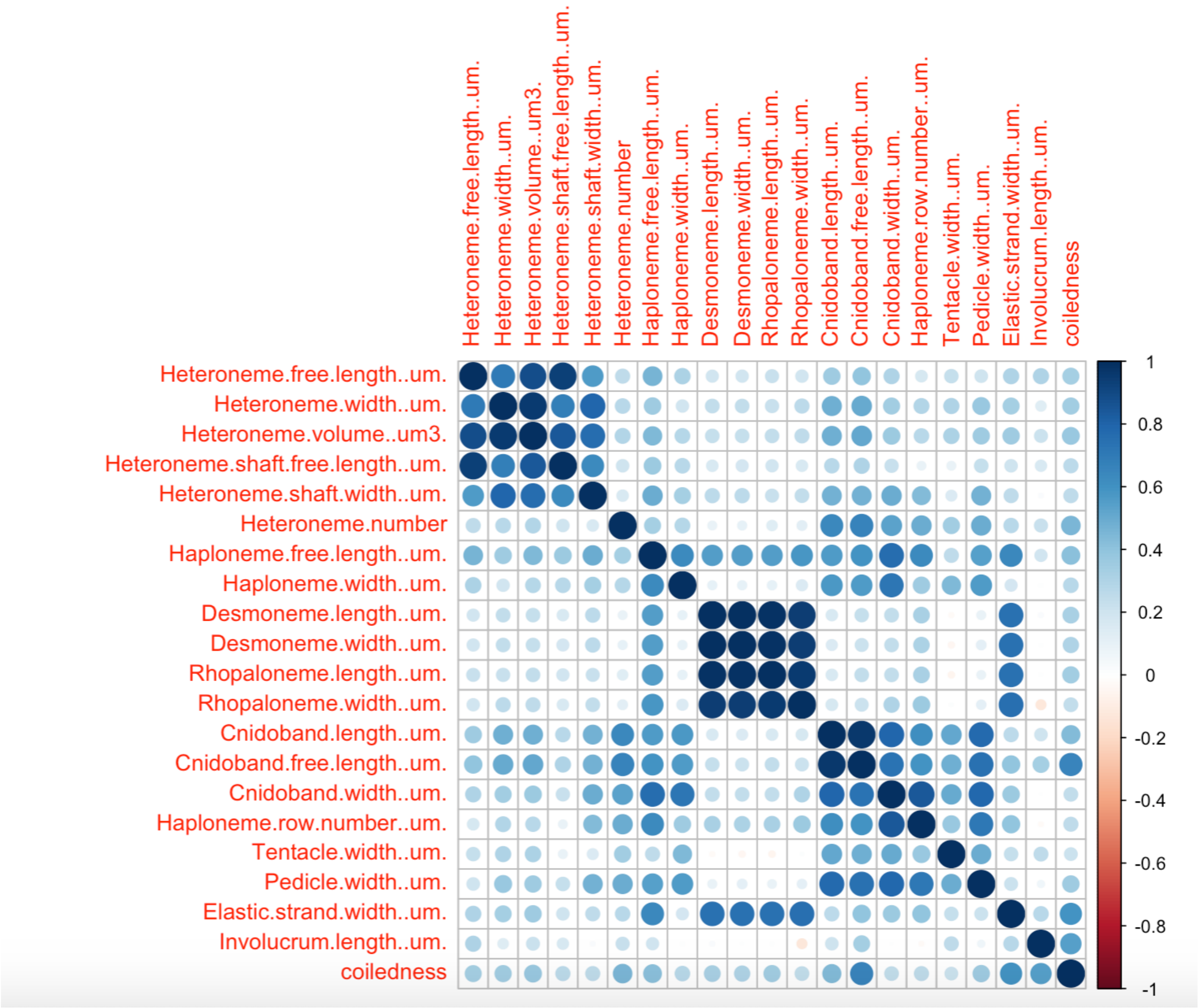


Figure 16: Rate covariance matrix for the whole tree using all taxa (45 species), transforming inapplicable states to zeroes. Covariances scaled to correlations. All characters estimated simultaneously under Brownian Motion.

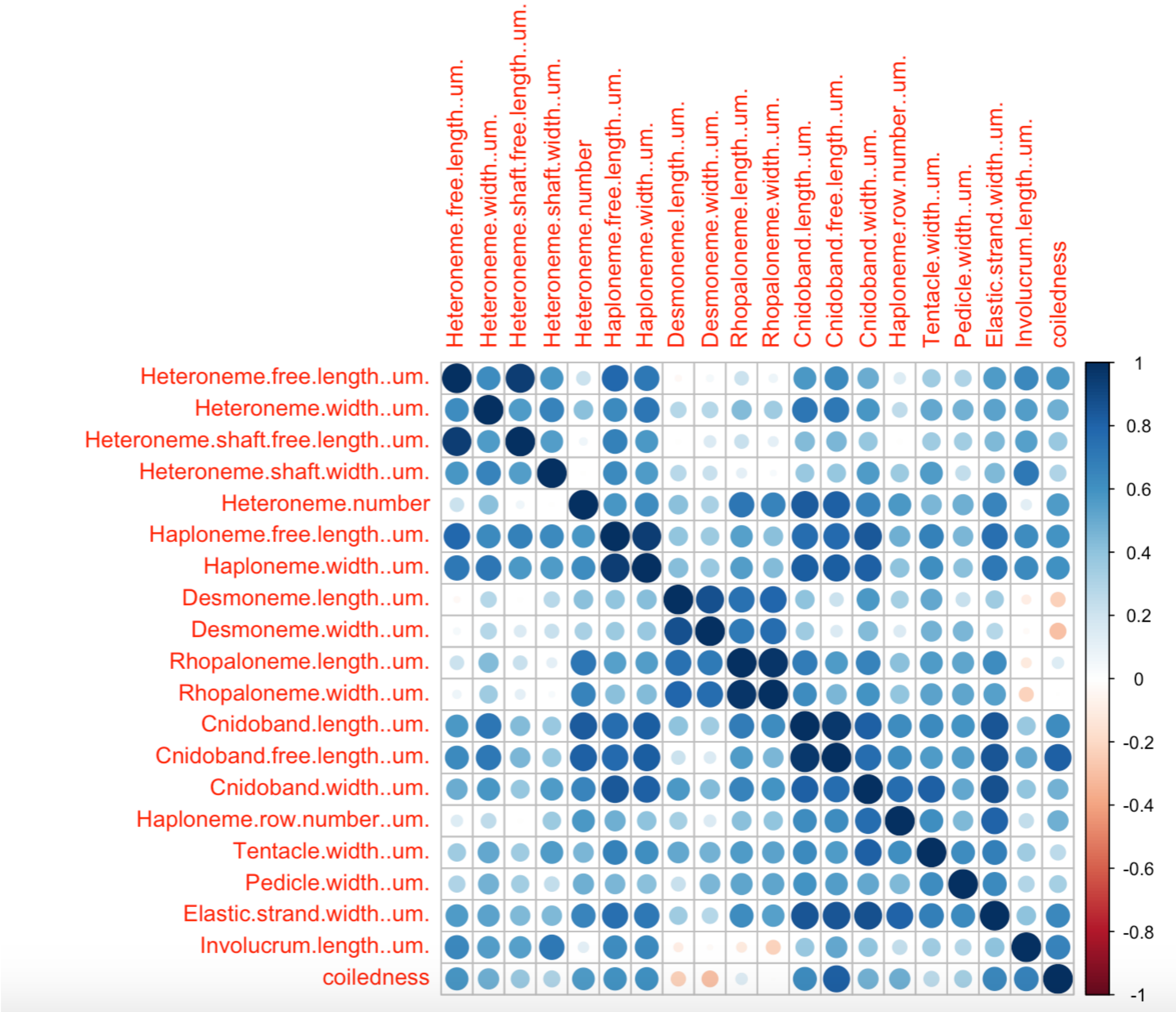


Figure 17: Rate covariance matrix for the whole tree using only taxa without inapplicable states (24 species). Covariances scaled to correlations. All characters estimated simultaneously under Brownian Motion.

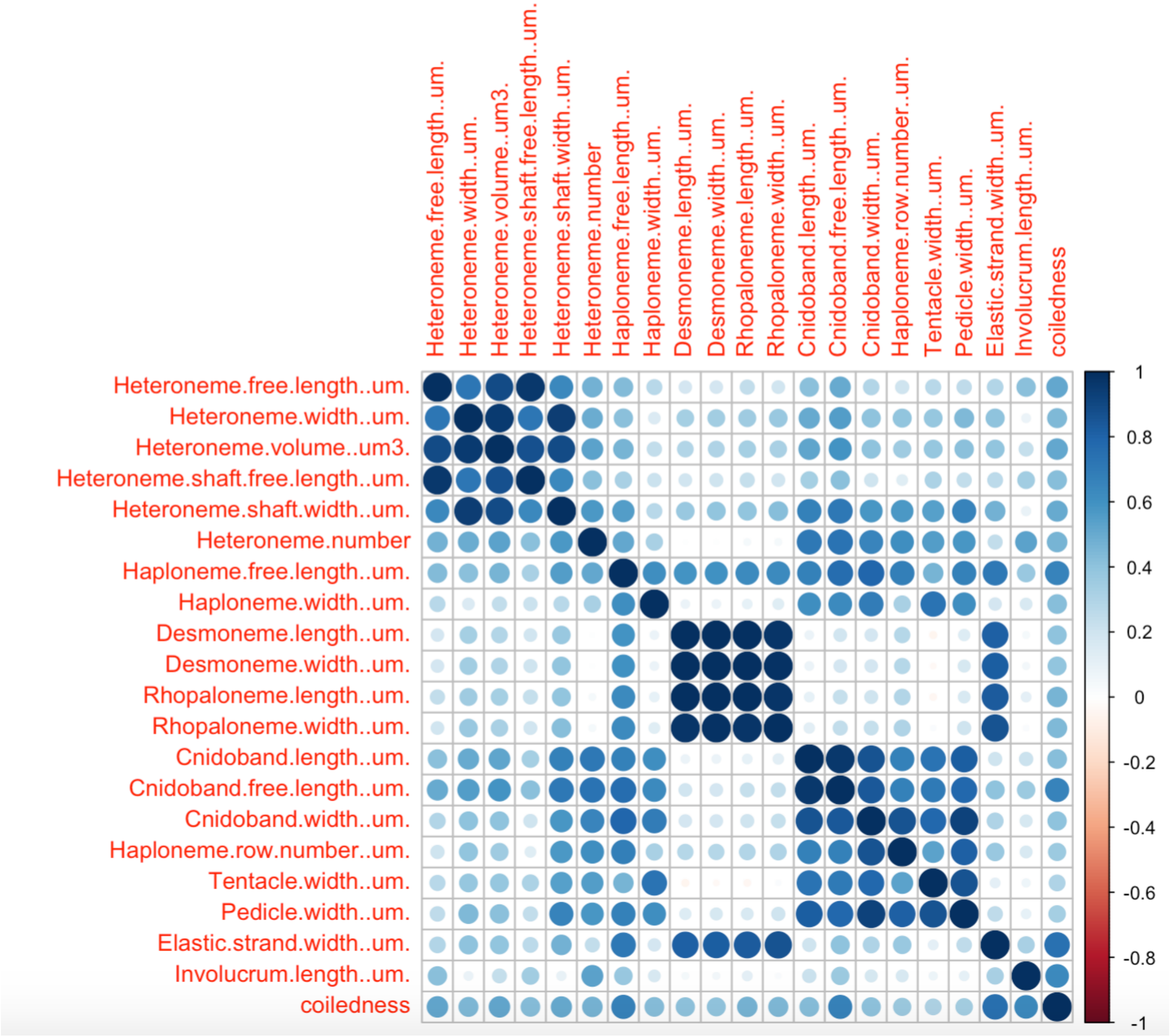


Figure 18: Rate covariance matrix for the whole tree using only taxa with diet data (22 species), transforming inapplicable states to zeroes. Covariances scaled to correlations. All characters estimated simultaneously under Brownian Motion.

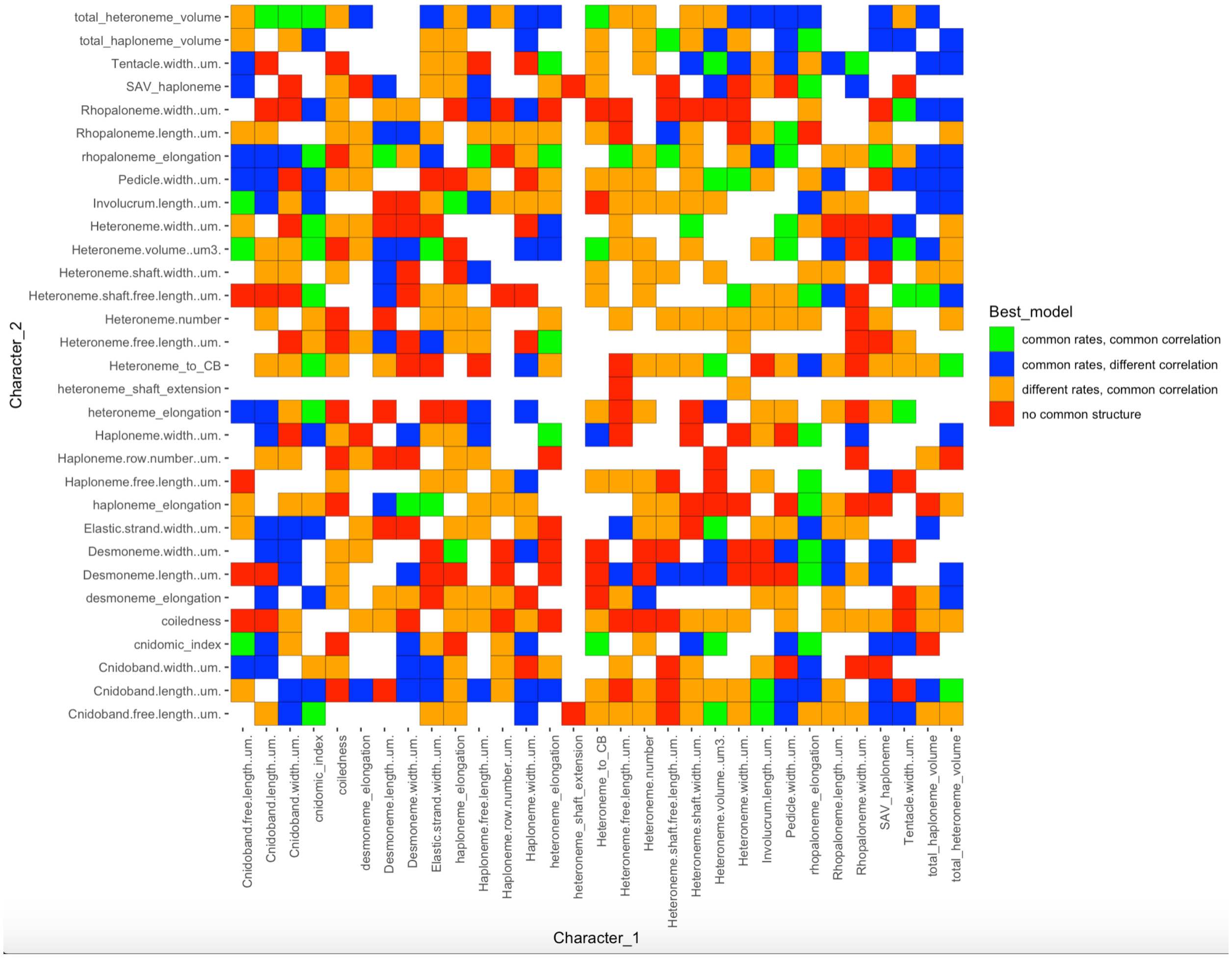


Figure 19: Best models (lowest AIC) supported in a pairwise character rate covariance analysis comparing correlated Brownian Motion models across the five selective regimes. Selective regimes were mapped onto the tree using an ancestral state reconstruction of the feeding guilds. Blank cells represent computationally singular contrasts.

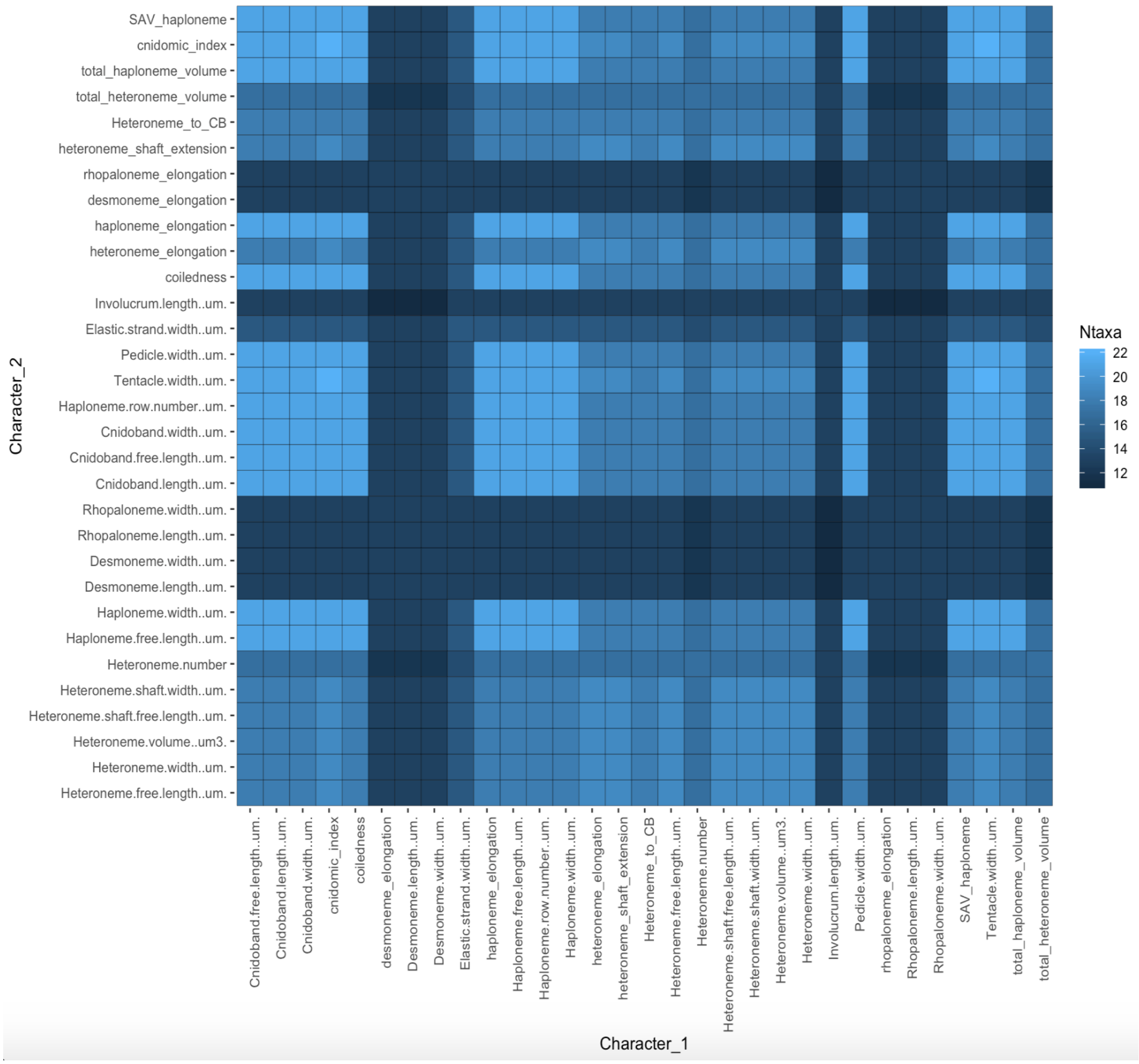


Figure 20: Number of taxa used for each pairwise contrast in the VCV analyses, given the number of taxa without inapplicable states.

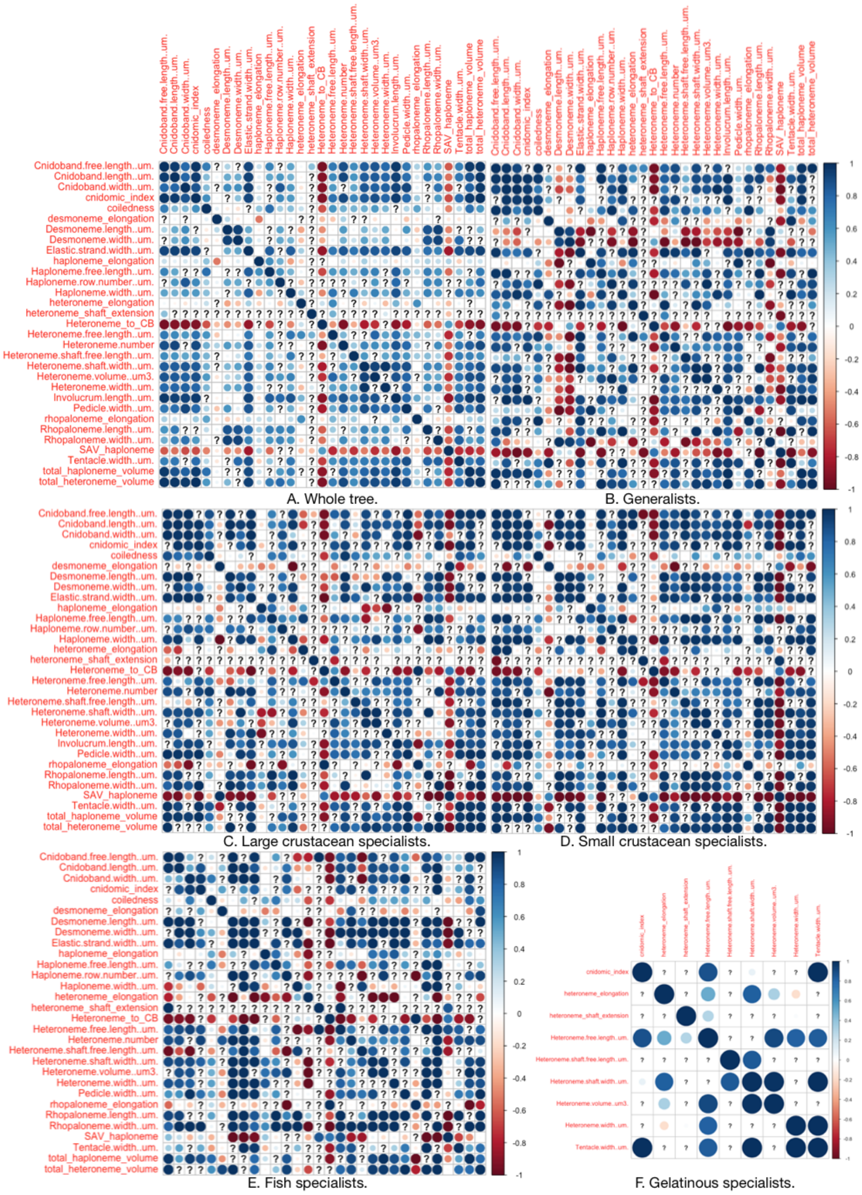


Figure 21: Pairwise estimated rate covariance matrices across the five selective regimes, using only taxa with diet data. Covariances scaled to correlations. Selective regimes were mapped onto the tree (22 species with diet data) using a stochastic mapping of the feeding guilds. Tree is pruned to taxa with no inapplicable states for a given character pair. Not all regimes are represented in all contrasts. Question marks represent computationally singular contrasts.

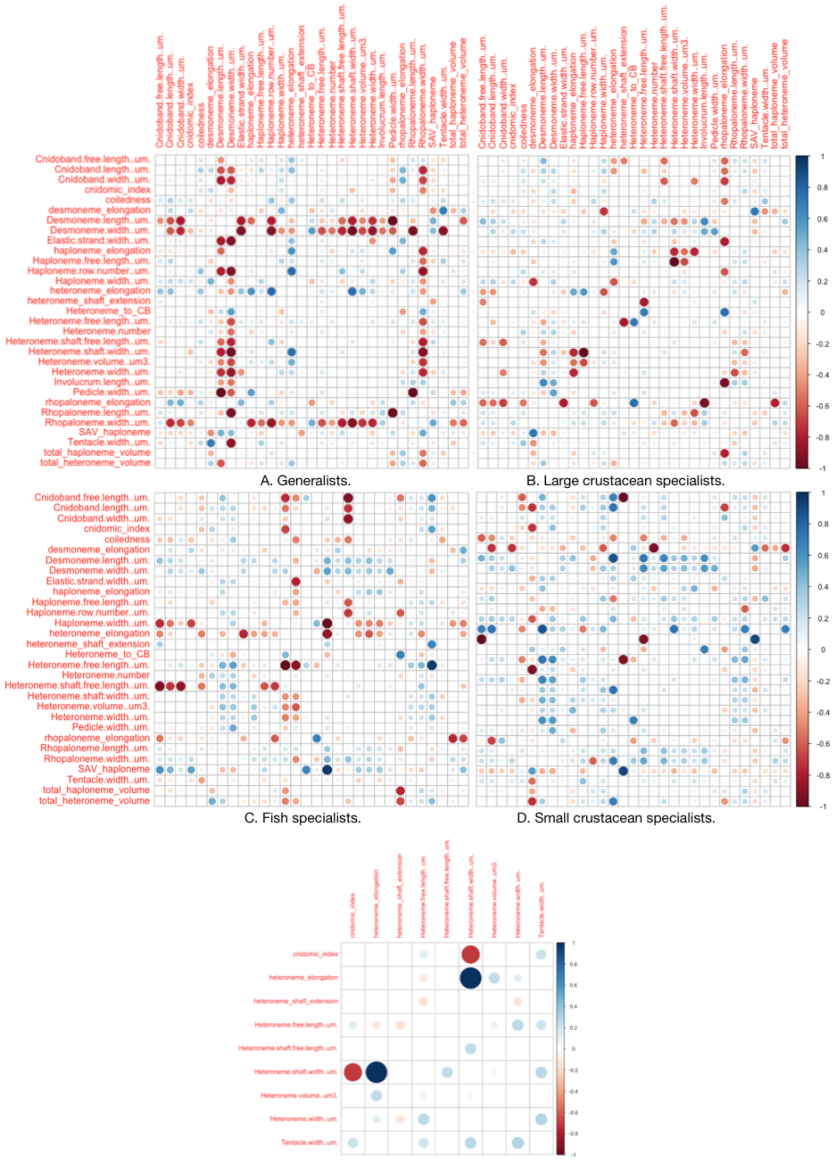


Figure 22: Scaled differences between the regime-specific covariance matrices in @ref{VCV_regimes} and the whole tree covariance matrix.

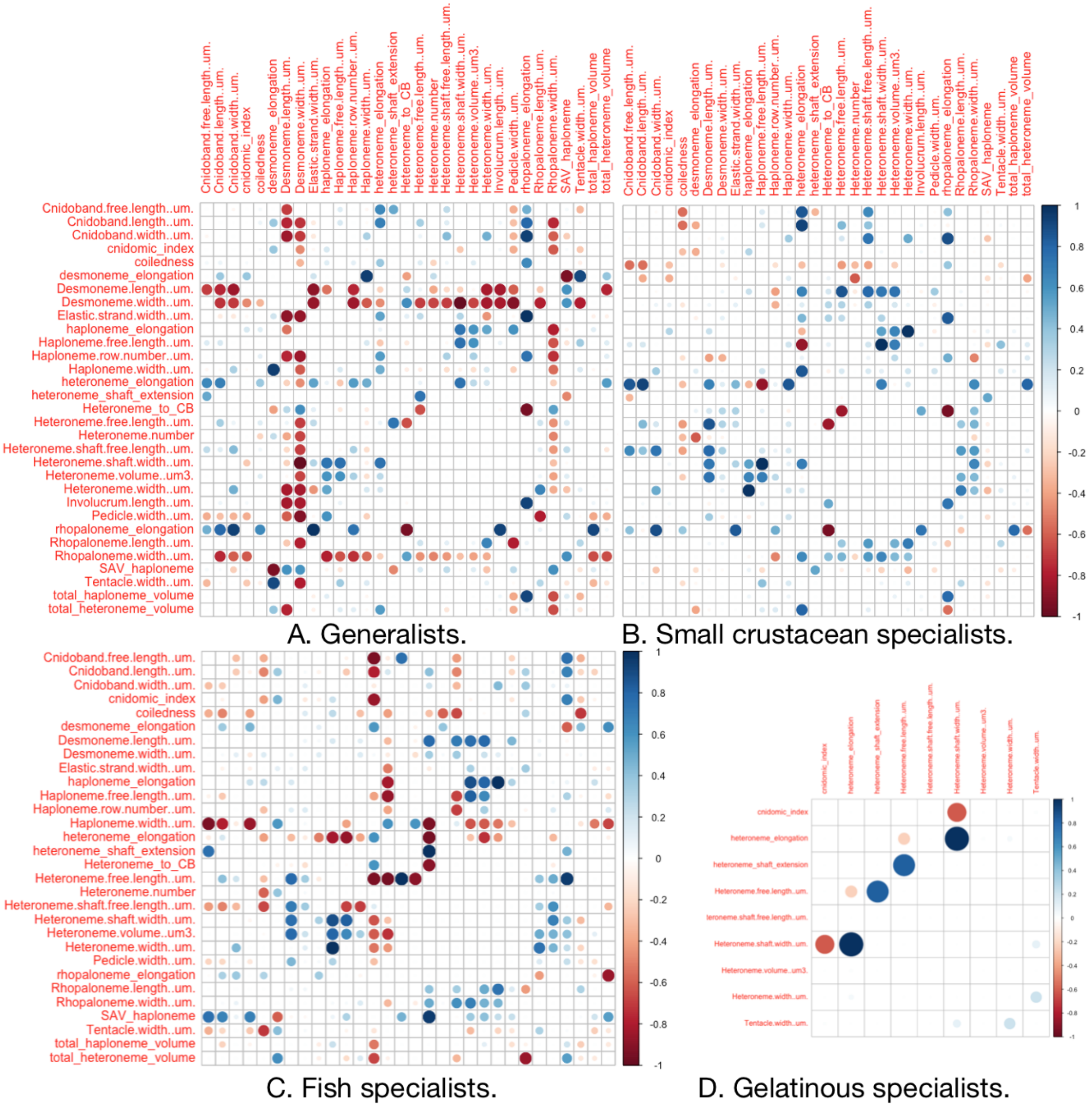


Figure 23: Scaled differences between the regime-specific covariance matrices in @ref{VCV_regimes} and the covariance matrices in their preceding regime, the large-crustacean specialist regime.
